# Nuisance effects in inter-scan functional connectivity estimates before and after nuisance regression

**DOI:** 10.1101/603183

**Authors:** Alican Nalci, Wenjing Luo, Thomas T. Liu

**Affiliations:** Center for Functional MRI, University of California San Diego, 9500 Gilman Drive MC 0677, La Jolla, CA 92093; Department of Electrical and Computer Engineering, University of California San Diego, 9500 Gilman Drive, La Jolla, CA 92093; Departments of Radiology, Psychiatry, and Bioengineering, University of California San Diego, 9500 Gilman Drive, La Jolla, CA 92093

**Keywords:** functional connectivity, nuisance regression, global signal, variability

## Abstract

In resting-state functional MRI, the correlation between blood-oxygenation-level-dependent (BOLD) signals across brain regions is used to estimate the functional connectivity (FC) of the brain. FC estimates are prone to the influence of nuisance factors including scanner-related artifacts and physiological modulations of the BOLD signal. Nuisance regression is widely performed to reduce the effect of nuisance factors on FC estimates on a per-scan basis. However, a dedicated analysis of nuisance effects on the variability of FC metrics across a collection of scans has been lacking. This work investigates the effects of nuisance factors on the variability of FC estimates across a collection of scans both before and after nuisance regression. Inter-scan variations in FC estimates are shown to be significantly correlated with the geometric norms of various nuisance terms, including head motion measurements, signals derived from white-matter and cerebrospinal regions, and the whole-brain global signal (GS) both before and after nuisance regression. In addition, it is shown that GS regression (GSR) can introduce GS norm-related fluctuations that are negatively correlated with inter-scan FC estimates. The empirical results are shown to be largely consistent with the predictions of a theoretical framework previously developed for the characterization of dynamic FC measures. This work shows that caution must be exercised when interpreting inter-scan FC measures across scans both before and after nuisance regression.

## 1. Introduction

Resting-state functional magnetic resonance imaging (fMRI) is a widely used method that aims to characterize the functional organization of the brain at rest (Smith et al., 2012; Fox and Raichle, 2007). The blood-oxygenation-level-dependent (BOLD) signal reflects metabolic changes in the brain that result from neuronal activity (Biswal et al., 1995; Hallquist et al., 2013). The correlation between BOLD signals across different brain regions is computed to estimate the functional connectivity (FC) of the brain (Raichle et al., 2001; Fox et al., 2005).

It is well-known that the BOLD signal is prone to the influence of various nuisance confounds including thermal noise, scanner drift, head motion, and physiological activity such as changes in respiration and heart rate (Bright and Murphy, 2015; Bright et al., 2017; Hallquist et al., 2013; Birn et al., 2008; Chang et al., 2009). If these confounds are not removed from the BOLD signal prior to analysis, they can lead to an increase in the number of false positives and negatives, causing erroneous interpretations of the fMRI results (Glasser et al., 2018).

Nuisance regression (NR) is widely performed to improve the spatial specificity of FC estimates on a per-scan basis. This involves projecting out a combination of nuisance measurements from the BOLD data prior to the computation of FC estimates. Nuisance measurements typically include but are not limited to head motion (HM) measurements, signals from the white-matter (WM) and cerebrospinal fluid (CSF) regions, cardiac and respiratory activity derived time courses, and the whole-brain global signal (GS) (Birn et al., 2008; Chang and Glover, 2009; Liu et al., 2015; Liu, 2016; Liu et al., 2017).

Despite the fact that NR is adopted with the assumption that it removes nuisance confounds from the FC estimates, it has been previously shown that it can be quiet ineffective in reducing the effects of nuisance confounds (Bright and Murphy, 2015; Power et al., 2012; Nalci et al., 2019). For example, HM regression has been shown to be a largely ineffective approach for reducing HM confounds in FC estimates even after projecting out 12 motion regressors (Power et al., 2012; Satterthwaite et al., 2012; Van Dijk et al., 2012).

More recently, in Nalci et al. (2019), we demonstrated that *dynamic* FC (DFC) estimates were related (within a scan) to the norms of various nuisance regressors such as the HM, WM+CSF, GS, heart rate, and respiratory derived time courses. We found that NR was largely ineffective in removing nuisance effects from DFC estimates with significant relations between the nuisance norms and DFC estimates remaining even after NR. We presented a theoretical framework to explain the limited effectiveness of NR and showed that the effects of nuisance norms on the DFC estimates were significant even when the correlations between the raw nuisance and BOLD signals were relatively small.

The fundamental difference between DFC and FC studies is the temporal duration over which the FC estimates (correlations between BOLD signals) are computed. In DFC studies the temporal window is typically on the order of 30-60 seconds, whereas in *static* FC studies the duration is the whole scan duration which is typically several minutes or longer (Hutchison et al., 2013; Preti et al., 2017). We will use the terms FC and static FC in an interchangeable fashion.

Although the effects of nuisance terms and efficacy of NR have been investigated on a per-scan basis (Liu et al., 2017), efforts to examine nuisance effects with regards to variations in FC estimates across scans have been rather limited. A better understanding of these effects is critical considering the increasing use of fMRI to examine the differences in FC measures between disease populations and healthy controls (Van Den Heuvel and Pol, 2010). Relevant studies include the investigation of FC metrics in Alzheimer’s disease (Greicius et al., 2004; Yang et al., 2014; Wang et al., 2007), Parkinson’s disease (Baudrexel et al., 2011), depression (Greicius et al., 2007), schizoprenia (Liu et al., 2008), dementia (Rombouts et al., 2009), and amyotrophic lateral sclerosis (ALS) (Mohammadi et al., 2009; Agosta et al., 2013).

In this work, we first investigate the effects of nuisance terms on the variability of FC estimates across different scans. Specifically, for each scan we compute the norms of the HM measurements, WM+CSF time courses, and the GS. We show the existence of significant correlations across scans between the FC estimates and each of the nuisance norms. We find nuisance regression using non-GS regressors to be largely ineffective in reducing the correlations between FC estimates and nuisance norms. We show that although GSR is partially effective in reducing the relation between GS norm and FC estimates, a considerable portion of the GS norm-related variance remains in the FC estimates, and strong GS norm-related fluctuations can be injected into the FC estimates.

Our work significantly extends the preliminary results regarding static FC estimates presented in Nalci et al. (2019). We provide a more extensive analysis of various nuisance effects on the FC estimates before and after NR. We generalize the theory developed in Nalci et al. (2019) to static FC measures and confirm the validity of the theoretical limitation of nuisance regression for correlation-based static FC estimates. We introduce *nuisance contamination maps* which illustrate the spatial distribution and extent of correlations between nuisance norms and FC estimates across scans. We also provide a detailed analysis of the limited efficacy of GSR and show how GSR can introduce GS norm-related fluctuations into the FC estimates.

## 2. Methods

### 2.1. Data

We used a publicly available dataset originally analyzed by Fox et al. (2007). The data were acquired from 17 young adults using a 3T Siemens Allegra MR scanner. Each subject underwent 4 BOLD echo-planar imaging (EPI) scans (32 slices, TR=2.16 s, TE=25 ms, 4×4×4 mm) each lasting 7 minutes (194 frames). The subjects were instructed to look at a cross-hair and asked to remain still and awake. High-resolution T1-weighted anatomical images were acquired for the purpose of anatomical registration (TR=2.1 s, TE=3.93 ms, flip angle=7 deg, 1×1×1.25 mm).

Standard pre-processing steps were conducted with the AFNI software package (Cox, 1996). The initial 9 frames from each EPI run were discarded to minimize longitudinal relaxation effects. Images were then slice-time corrected and co-registered, and the 6 head motion parameter time series were retained. The images were converted to Talairach and Tournoux (TT) coordinates, resampled to 3 mm cubic voxels, and spatially smoothed using a 6 mm full-width-at-half-maximum isotropic Gaussian kernel. The 0^*th*^ and 1^*st*^ order Legendre polynomials (a constant term to model the temporal mean and a linear trend) were projected out from each voxel’s time course. Each voxel time series was then converted into a percent change BOLD time series through division by the estimate of the temporal mean. This version of the data will be referred to as “uncorrected” data in this paper.

We used seed signals derived from the posterior cingulate cortex (PCC), intraparietal sulcus (IPS), frontal eye fields (FEF), auditory (AUD) and motor (MOT) networks. These seed signals were obtained by averaging time series selected over spheres of radius 6 mm (2 voxels) centered about their corresponding TT coordinates (He and Liu, 2012). The sphere centers were obtained by converting the MNI coordinates from Van Dijk et al. (2010) to TT coordinates (Lacadie et al., 2008). For the PCC, IPS, FEF and MPF seeds we used the coordinates [0,-51,26], [32,-51,41], [24,-13,51], and [6,32,28], respectively. For the left MOT, and right MOT seeds we used the coordinates [-36,-22,52] and [37,-12,52], respectively. A combined MOT seed was obtained by using the left and right MOT coordinates to define two spheres and by merging the spheres. A combined AUD seed was obtained by using the left and right AUD coordinates [-41,-26,14] and [41,-26,14], respectively. Finally, for the WM and CSF nuisance signals we defined the sphere centers as [31,-28,32] and [-15,-28,21], respectively.

### 2.2. Inter-scan variations in FC estimates

To investigate the variations in FC estimates across scans, we computed the Pearson correlation between a seed signal and every other voxel in the brain for each scan. Denoting the zero mean percent change BOLD signals from a seed-voxel pair as *x*_1_ and *x*_2_ in vector notation, the FC estimate for the *k*th scan was obtained by computing 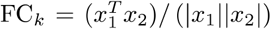, where |.| denotes the *ℓ*_2_ norm and *k* is the scan index. For each seed-voxel pair, we computed the FC estimates across all scans and concatenated them to form a vector of FC estimates: FC_Vec_ = {FC_1_, FC_2_,…, FC_*K*_} where *K* = 68 is the total number of scans. This vector will be referred to as the inter-scan FC estimates or simply as FC estimates. We obtained a separate vector FC_Vec_ for each seed voxel pair in the brain (i.e. for a single seed, we have *N* vectors where *N* is the number of voxels).

### 2.3. Nuisance regressions

To investigate the effects of nuisance regression on FC estimates, we performed 4 separate nuisance regressions on the uncorrected data. This was done prior to the computation of FC estimates. Nuisance regressions involved projecting out (1)6 HM parameters, (2) 6 HM parameters combined with the signals from the WM and CSF regions, (3) the GS time course, and (4) HM, WM, CSF signals combined with the GS. The global signal (GS) was obtained as the average of all (percent) change BOLD time courses across the whole brain volume. For each nuisance regression, the vector of inter-scan FC estimates prior to nuisance regression will be referred to as “Pre FC” estimates and after regression as “Post FC” estimates.

### 2.4. Norm as a nuisance metric on FC estimates across scans

To measure the effect of nuisance terms on the FC estimates across scans we adopted the approach in Nalci et al. (2019). For GS regression, we first computed the *ℓ*_2_ norm of the GS time course for each scan. Denoting the GS time course for a scan *k* with *n_k_*, we computed the *ℓ*_2_ norm as 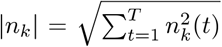, where *t* indexes over time and *T* is the total number of time points. We then concatenated the GS norms across different scans to obtain a vector of GS norms as |*n*|_Vec_ = {|*n*_1_|, |*n*_2_|,…,|*n_K_*|}.

For multiple nuisance regressions (e.g. HM+WM+CSF) we obtained a total norm of all the regressors involved by computing 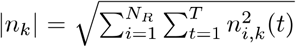, where *n_i,k_* is a single regressor time course, *i* is the index over multiple regressors and *N_R_* is the total number of regressors. Finally, we concatenated the corresponding total nuisance norms across different scans to obtain a nuisance norm vector as |*n*|_Vec_ = {|*n*_1_|,|*n*_2_|,…,|*n_K_*|}.

### 2.5. Nuisance contamination maps: Nuisance contamination of FC estimates across scans

We quantify the nuisance contamination in inter-scan FC estimates by correlating the nuisance norm vectors |*n*|_Vec_ with the vector of FC estimates FC_Vec_ for each seed-voxel pair. This approach is illustrated in Figure 1. In the top row, we first computed the correlations between a seed signal (e.g. PCC seed shown with red color) and the time series from every other voxel (lines with blue color) to form a seed-based correlation map for each scan (represented as a *N* × 1 column vector with red color). We then repeated this for all 68 scans and concatenated the resulting seed-based correlation vectors (FC maps) to form a (*N* × 68) matrix as shown in the left hand side in the second row. Each row of this matrix corresponds to the FC estimates vector for a single seed voxel pair (an example row is shown with green color). We then computed the correlations between the FC estimates rows and nuisance norm vector (time series with black color) to form a nuisance contamination vector which can be reshaped into a 3D nuisance contamination map. The green colored square in the nuisance contamination vector corresponds to a single correlation coefficient obtained between the nuisance norm and the FC vector from a single seed voxel pair. This is also depicted on the nuisance contamination map with the green (+) symbol.

**Figure 1:**
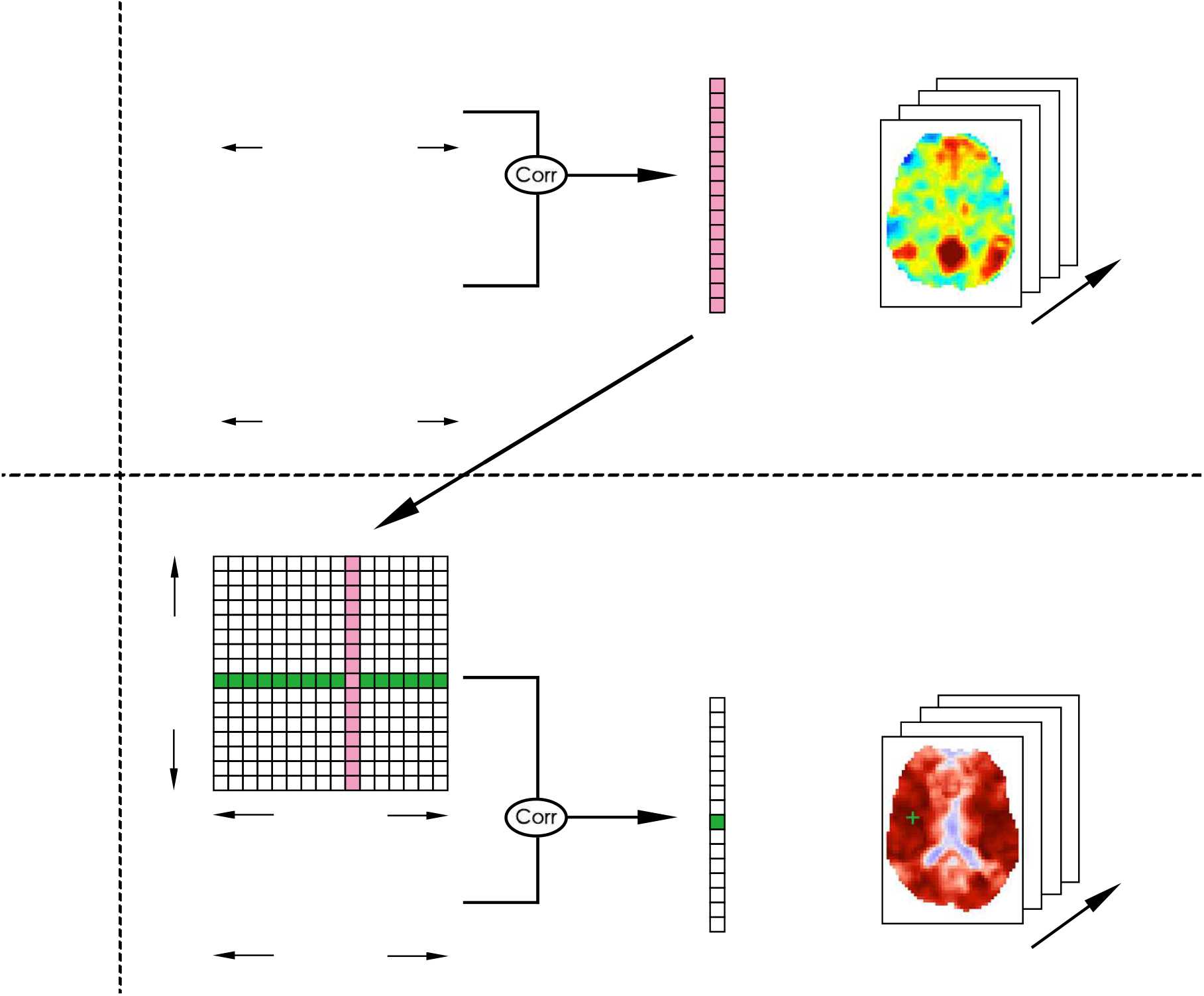
Diagram illustrating how to obtain nuisance contamination maps. These maps visualize the spatial distribution of the correlations between inter-scan FC estimates (FC_Vec_) and nuisance norms (|*n*|_Vec_). In the first row, a seed signal (e.g. PCC seed) is correlated with signals from other voxels in the brain to form a vector (vertical vector with red color) of FC estimates. This vector represents the canonical seed-based PCC correlation map. This step is repeated across all 68 scans and the resulting PCC maps are concatenated column-wise to form a matrix of FC estimates across scans. This matrix is shown on the bottom left-hand side. The individual rows of the matrix correspond to inter-scan variations in the FC between the PCC seed and a single voxel (an example is shown with green color). Each column is the PCC map for a single scan. The individual rows of this matrix are then correlated with the nuisance norm (shown at the bottom left) to obtain the nuisance contamination values as a 1D vector. An entry of the nuisance contamination vector corresponds to the correlation between the nuisance norm and the FC vector from a single seed-voxel pair. This is illustrated with the dark green (+) symbol on the contamination map. The nuisance contamination vector can be reshaped into a 3D volume to investigate regions of nuisance contamination across the brain.

We obtained nuisance contamination maps both before and after each regression and for different seed signals including the PCC, IPS, FEF, MOT, and AUD seeds. Note that these maps are **not** functional connectivity maps, but instead quantify the relations between seed-based FC estimates and nuisance norms across different scans.

### 2.6. Theoretical bound on ΔFC

In Nalci et al. (2019) we presented a theoretical expression for the difference ΔDFC = (Post DFC – Pre DFC) between the dynamic FC (DFC) estimates obtained before and after nuisance regression in seed correlation-based DFC studies. This mathematical theory applies for static FC studies as well by noting that a temporal sliding window in DFC analysis can be replaced by the whole scan duration. Thus, the following theoretical bounds apply for the difference between Pre FC and Post FC estimates obtained before and after nuisance regression:

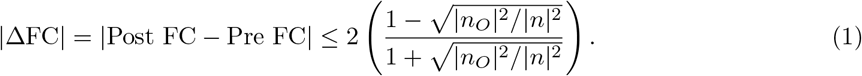

Here, *n* is a single nuisance regressor time course represented in vector notation. The nuisance regressor can be decomposed as *n* = *n_I_* + *n_O_*, where *n_I_* is an in-plane component that lies in the subspace spanned by a single seed-voxel pair *x*_1_ and *x*_2_ and *n_O_* is the component orthogonal to this subspace.

The orthogonal nuisance fraction 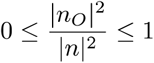 reflects the nuisance energy that lies in the orthogonal subspace and serves as a measure of orthogonality between the nuisance regressor and the seed-voxel pair (e.g. *x*_1_ and *x*_2_). If *n_O_* becomes arbitrarily large then the fraction 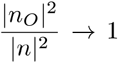 and |ΔFC| → 0. An example of this bound is provided in Figure 5 where a large orthogonal nuisance fraction for the HM, WM, CSF and HM+WM+CSF regressors impose a narrow bound on |ΔFC| values forcing them to cluster close to 0.

Note that an exact value for the orthogonal nuisance fraction 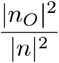 can be obtained when using a single regressor such as the GS. In the case of multiple regressors, an estimate of the orthogonal nuisance fraction and |ΔFC| can be obtained by using the first principal component (PC) of the multiple regressors as in Nalci et al. (2019). This simple approximation enables us to understand the approximate relation between |ΔFC| and the orthogonal nuisance fraction. When we analyze multiple regressors, we will provide the approximate orthogonal nuisance fraction values and will also show that regression with the first PC is a good approximation to performing multiple regression.

### 2.7. Significance testing of the relation between FC variations and nuisance norms across scans

We assessed the statistical significance of the relation between the FC estimates and nuisance norms across scans using non-parametric null testing. As the ordering of scans is not important, we formed null distributions by randomly permuting the scan ordering of FC estimates for each seed voxel pair and nuisance norm over 10,000 trials. We then correlated the resulting surrogate FC estimates with nuisance norms and obtained 10,000 null correlation values both before and after nuisance regression. We used the null distributions to assess the statistical significance of the correlations between the non-permuted FC estimates and nuisance norms.

## 3. Results

In this section we show that variations in FC estimates across multiple scans are significantly correlated with the geometric norms of various nuisance terms. We demonstrate that a considerable portion of the FC estimates remains significantly correlated with nuisance norms even after nuisance regression. We make use of the theoretical findings from Nalci et al. (2019) to show that the inefficacy of nuisance regression for non-GS regressors such as HM, WM and CSF is largely due to the large orthogonality between nuisance regressors and the BOLD data within each scan. We further show that GSR can introduce negative GS norm fluctuations into the FC estimates.

In the first row of Figure 2 we show examples of the relation between nuisance norms and FC estimates for 3 seed pairs: PCC&AUD, MPF&AUD and PCC&IPS. The column labels indicate both the type of nuisance norm and the specific seed-pair. The FC estimates in each column (blue lines, labeled as Pre FC) are significantly (*p* < 10^−3^) correlated with various nuisance norms (black lines) before nuisance regression. The correlations obtained between the Pre FC estimates and HM, HM+WM+CSF, and GS norms in the first, second, and third columns are *r* = 0.56, *r* = 0.58, and *r* = 0.82, respectively. After nuisance regression, the Post FC estimates are still significantly correlated with the nuisance norms with correlations of *r* = 0.50, *r* = 0.55, and *r* = 0.54 observed for HM, HM+WM+CSF, and GS norms, respectively. The second row shows the same relations using scatter-plots where the FC estimates are plotted against the respective nuisance norms.

**Figure 2:**
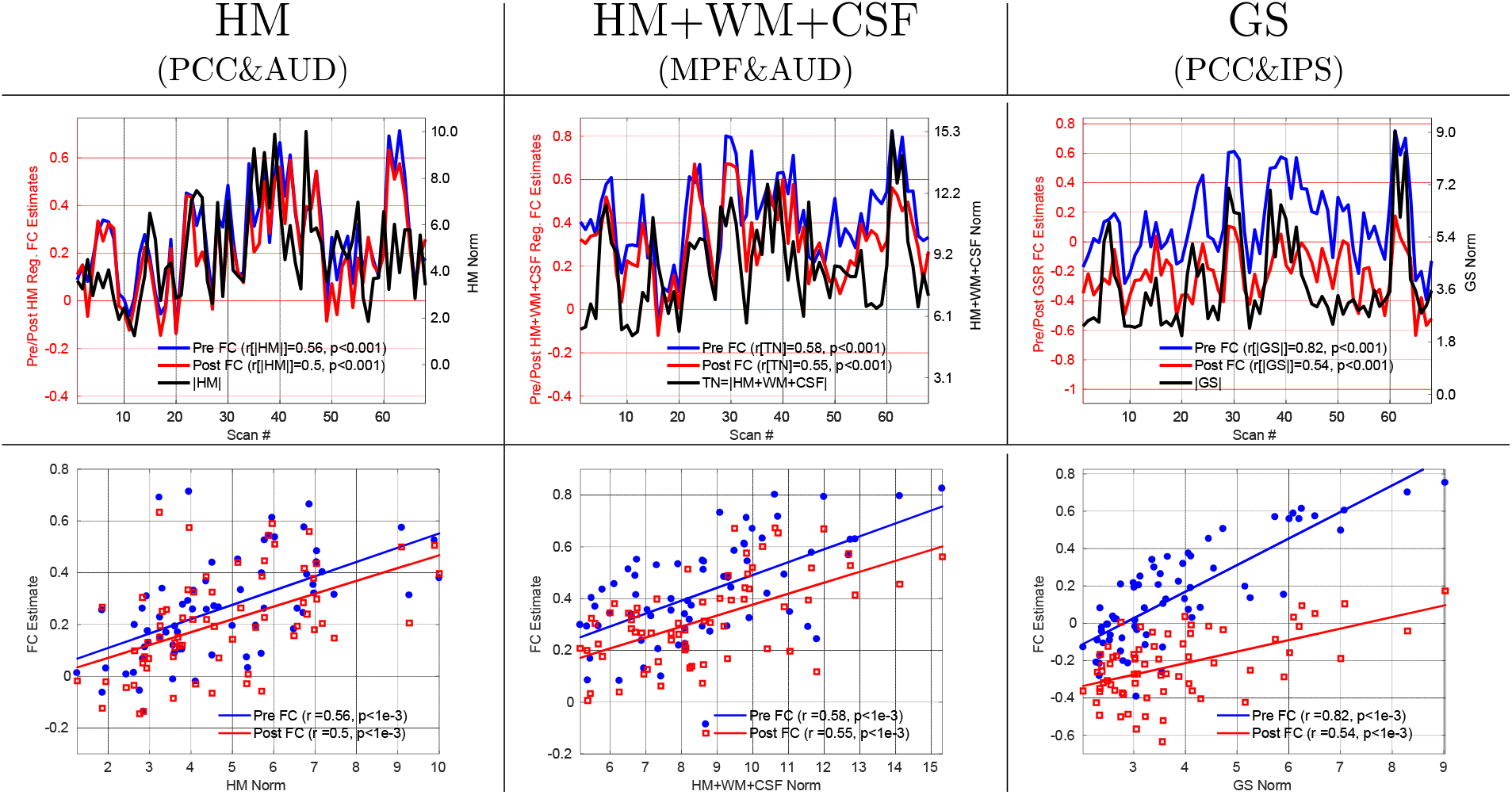
FC estimates obtained from 3 example seed pairs both before (blue lines) and after (red lines) nuisance regression are significantly correlated (*p* < 10^−3^) across scans with various nuisance norms (black lines). The type of nuisance norm and seed pair are indicated by the column labels. In the first row the FC estimate values are indicated on the left y-axis in each column, nuisance norm values are indicated on the right y-axis, and scan numbers are indicated by the x-axis. Before nuisance regression, the correlations between the Pre FC estimates and nuisance norms were *r* = 0.56, *r* = 0.58, and *r* = 0.82 for the HM, HM+WM+CSF and GS norms, respectively. After nuisance regression, the correlations between the Post FC estimates and nuisance norms were *r* = 0.50, *r* = 0.55, and *r* = 0.54 for the HM, HM+WM+CSF and GS norms, respectively. The first row serves as a nice visual demonstration of the similarity between the fluctuations in nuisance norms and FC estimates. The second row shows the relation between the FC estimates and nuisance norms using scatter-plots.

The results presented below generalize the relation between various nuisance norms and PCC-based FC estimates to include all PCC seed-voxel pairs. We provide the results for other seeds in the supplementary material and main text below.

### 3.1. HM regression

In Figure 3a,b we show the PCC-based HM contamination maps before and after HM regression. These maps are very similar to each other (cosine similarity *S* = 0.98) and show widespread correlations between the HM norm and FC estimates across scans both before and after HM regression.

**Figure 3:**
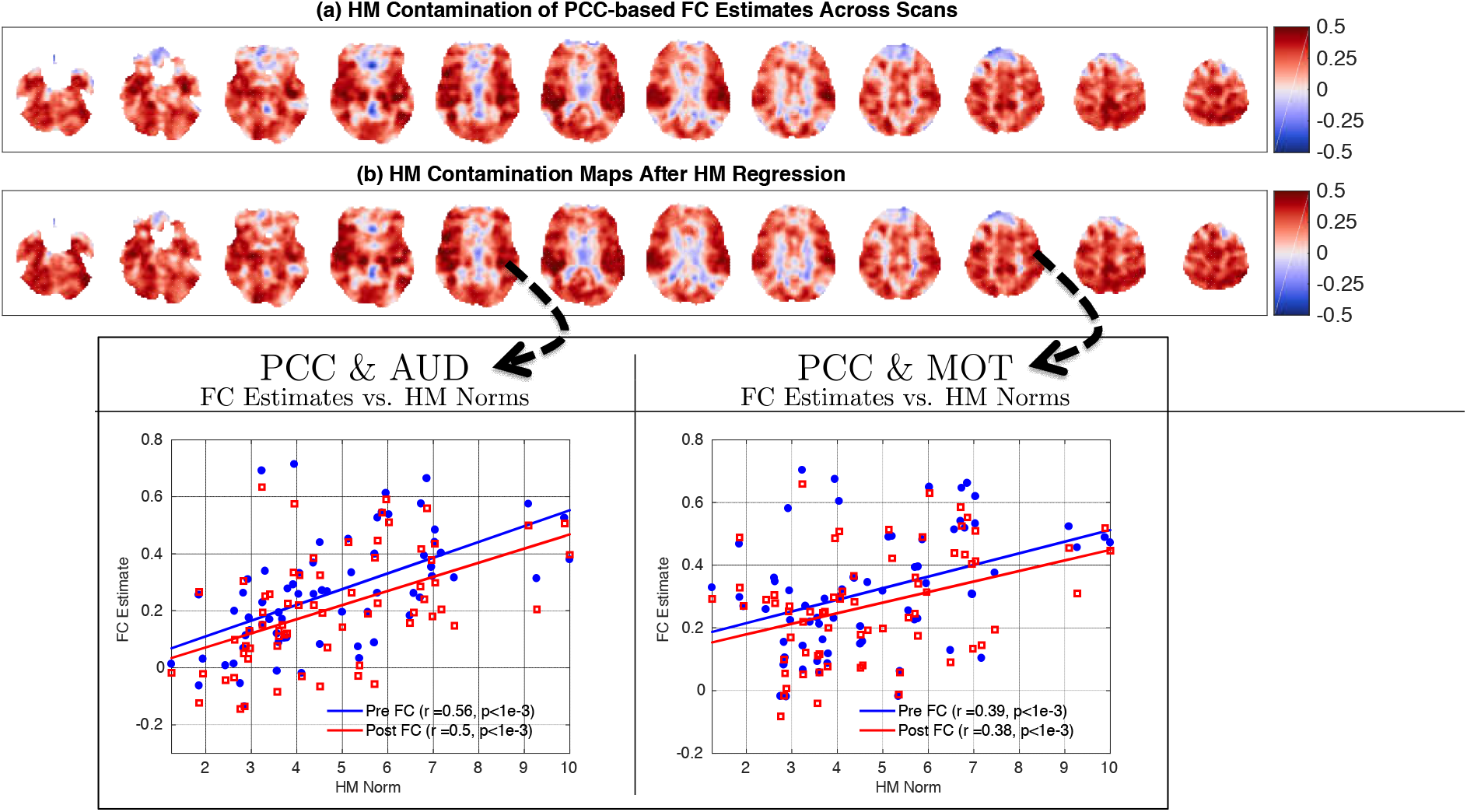
HM contamination maps obtained by correlating the HM norm with FC estimates (a) before and (b) after HM regression. The contamination maps in (a) and (b) are fairly similar to each other (cosine similarity *S* = 0.98) and both show widespread correlations between the HM norm and FC estimates across scans. This indicates that HM regression is largely ineffective in removing the relation between the HM norm and FC estimates. We show two example seed-pairs (PCC & AUD and PCC & MOT) at the bottom to illustrate the relation between HM norms and FC estimates for two different regions.

In the scatter plot shown in Figure 4 we plot the correlations obtained between the Post FC estimates and HM norm versus the correlations obtained between the Pre FC estimates and HM norm. The sideways histogram along the y-axis shows the distribution of correlation values obtained between the Post FC estimates and HM norm, which ranged from *r* = −0.36 to *r* = 0.60 with a mean of 0.27. The histogram along the x-axis at the bottom shows the correlations obtained between the Pre FC estimates and HM norm, which ranged from *r* = −0.34 to *r* = 0.61 with a mean of 0.26. These two correlation distributions were strongly related to each other (*r* = 0.94, *p* < 10^−3^). The linear fit (blue line) between the correlation distributions (Slope= 0.91, Offset= 0.04) was very close to the line of unity (yellow dashed line). This indicates that HM regression had a very limited effect in removing the correlations between the Pre FC estimates and HM norms, and the correlations with the HM norm largely persisted after HM regression.

**Figure 4:**
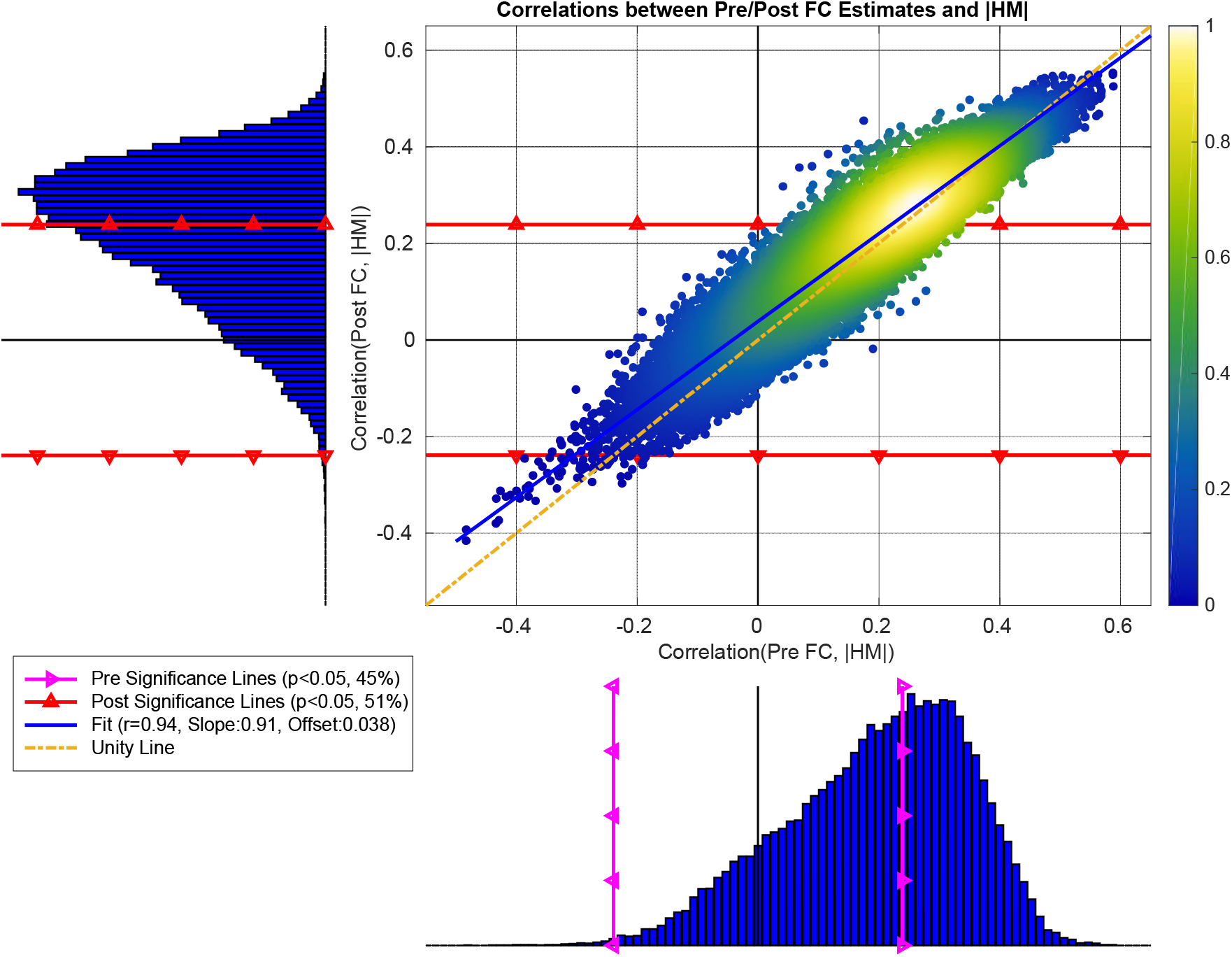
The correlations between the Post FC estimates and HM norm versus the correlations obtained between Pre FC estimates and HM norm. These correlation distributions were significantly related (*r* = 0.94, *p* < 10^−6^) to each other. The linear fit (blue line, Slope= 0.91 and Offset= 0.04) between the two correlation distributions was fairly close to the line of unity (dashed yellow line). At the bottom histogram, the correlations between the Pre FC estimates and HM norms ranged from *r* = −0.34 to *r* = 0.61 with mean 0.26. We superimposed the significance lines (*p* < 0.05) on this histogram using magenta lines with triangles (labeled as Pre Significance). Based on the significance line, 45% of the Pre FC estimates were significantly correlated with HM norms. In the sideways histogram to the left, the correlations between the Post FC estimates and HM norms ranged from *r* = −0.36 to *r* = 0.60 with mean 0.27, where significance lines are shown with red lines with triangles. 51% of the Post FC estimates were significantly correlated with HM norms after HM regression.

As shown in Table 1, 45% of the PCC-based Pre FC estimates were significantly correlated (*p* < 0.05) with HM norms. We further computed the mean and standard deviation of the percent variance explained by the HM norm for those significant 45% Pre FC estimates. The HM norm explained an average of 11%±4% (mean+SD) of the variance in those significant Pre FC estimates. After HM regression, 51% of the Post FC estimates were significantly correlated with the HM norm. The HM norm explained an average of 11%±4% (mean+SD) of the variance in those Post FC estimates, which is the same percent variance explained prior to HM regression.

**Table 1:** Summary of the relationship between the FC estimates and nuisance norms obtained before and after nuisance regressions for 5 seed signals. The seed signal and regression state (e.g. before and after) are indicated in the first and second columns, respectively. For each nuisance regression we report the percentage of seed-voxel pairs for which there was a significant correlation between the FC estimates and nuisance norms. We also provide the mean and standard deviation of the percent variance explained in the FC estimates by the nuisance norms over the respective subsets of significant seed-voxel pairs.

| Seed | State | HM |  | HM+WM+CSF |  | GS |  | HM+WM+CSF+GS |  |
| --- | --- | --- | --- | --- | --- | --- | --- | --- | --- |
| | | %Brain | %Var<br>mean±SD | %Brain | %Var<br>mean±SD | %Brain | %Var<br>mean±SD | %Brain | %Var<br>mean±SD |
| PCC | Pre FC | 45 | 11±4 | 88 | 27±12 | 96 | 37±15 | 91 | 31±14 |
| | Post FC | 51 | 11±4 | 56 | 13±6 | 27 | 11±5 | 26 | 11±5 |
| MOT | Pre FC | 45 | 13±5 | 96 | 26±9 | 99 | 38±13 | 98 | 30±10 |
| | Post FC | 49 | 13±5 | 74 | 15±7 | 30 | 13±6 | 20 | 11±5 |
| AUD | Pre FC | 60 | 13±5 | 98 | 28±10 | 99 | 40±13 | 98 | 33±12 |
| | Post FC | 63 | 14±6 | 75 | 16±8 | 25 | 11±5 | 22 | 10±4 |
| FEF | Pre FC | 43 | 11±4 | 100 | 32±9 | 99 | 49±11 | 99 | 38±10 |
| | Post FC | 49 | 11±4 | 82 | 17±8 | 24 | 11±5 | 23 | 11±5 |
| IPS | Pre FC | 40 | 11±4 | 97 | 25±9 | 99 | 40±13 | 98 | 29±10 |
| | Post FC | 43 | 11±4 | 63 | 13±6 | 20 | 12±5 | 23 | 11±5 |

Ideally, HM regression should fully remove the HM norm variance from the Post FC estimates. However, from Figure 4 we see that the Post FC estimates are still correlated with HM norm nearly as much as the Pre FC estimates are correlated with HM norm. In Figure 5a we show the theoretical limitation of HM regression. Each point in this plot shows the ΔFC (difference between the Pre FC and Post FC estimates) versus the orthogonal nuisance fraction |*n_O_*|^2^/|*n*|^2^ for a single seed-voxel pair in a single scan. The HM regressors are largely orthogonal to most seed voxel pairs across scans with a mean orthogonal fraction of |*n_O_*|^2^/|*n*|^2^ = 0.99. Due to the large orthogonal fraction, the theoretical bounds force the ΔFC values to be clustered around a mean value of 0. Since HM regression cannot really alter the Pre FC estimates, the slope of the linear fit in Figure 4 remains close to the line of unity.

**Figure 5:**
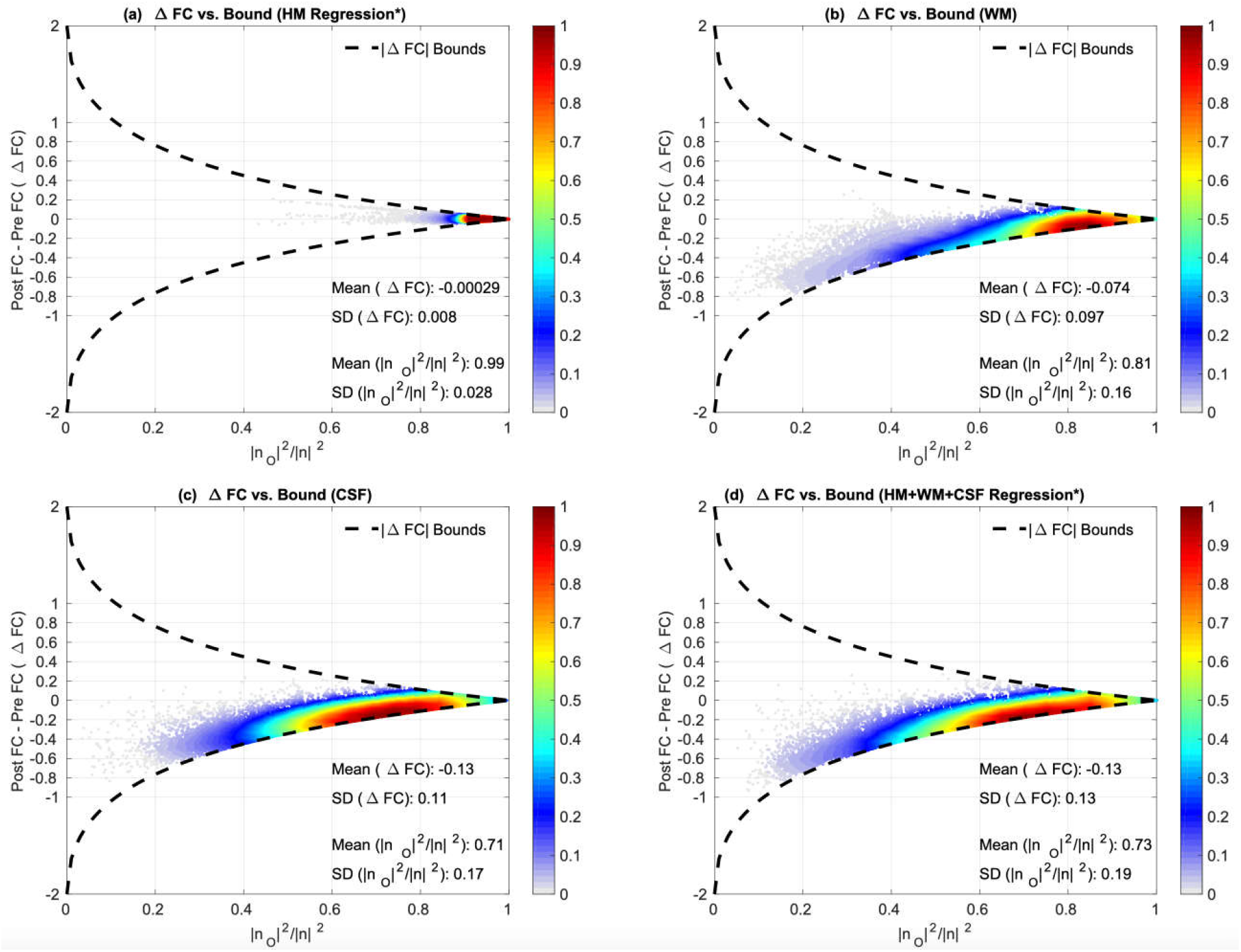
ΔFC versus orthogonal nuisance fraction 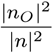 for HM (*1^st^ PC), WM, CSF and HM+WM+CSF (*1^st^ PC) regressor groups. Each point in these plots represent the values for a single scan and a single seed-voxel pair. In (a) we plot the ΔFC versus the orthogonal nuisance fraction for HM regression. We see that the points are strictly clustered on the right hand side around a mean orthogonal nuisance fraction value of 0.99. This imposes an extremely tight bound on the ΔFC values before and after regression and HM regression resulting in a negligible effect on the FC estimates (mean ΔFC = 0). In (b), (c) and (d) we plot the ΔFC versus the orthogonal nuisance fraction for the WM, CSF, and 1^st^ PC of HM+WM+CSF regressors, respectively. These regressors are largely orthogonal to BOLD data with mean orthogonal fractions 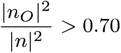, resulting in fairly tight bounds on the ΔFC values. Thus, the effects of these regressors on the FC estimates are very limited with mean |ΔFC| ≤ 0.13.

Note that as in Nalci et al. (2019) we used the first principal component (PC) across all head motion measurements for the ΔFC plot in Figure 5a. We provide the ΔFC plot after performing multiple HM regression in Supplementary Figure 1, which shows that ΔFC values are still largely clustered around zero (mean ΔFC = −0.037), roughly within the tight theoretical bounds.

We present the HM contamination maps and their respective correlation distributions before and after HM regression for other seeds (e.g. MOT, AUD, FEF, and IPS) in Supplementary Figures 3 to 6. These were similar to the PCC-based results discussed above. Additionally, Table 1 summarizes the significance results of the relation between nuisance norms and FC estimates for other seeds.

### 3.2. HM+WM+CSF regression

In Figure 6a,b we show the HM+WM+CSF contamination maps both before and after nuisance regression. These maps show widespread correlations between the nuisance norm and FC estimates across scans. In panel (b) there is a visible reduction in the positive correlation values (i.e. red regions) after regression with a slight increase in anti-correlations (i.e. blue regions start to appear).

**Figure 6:**
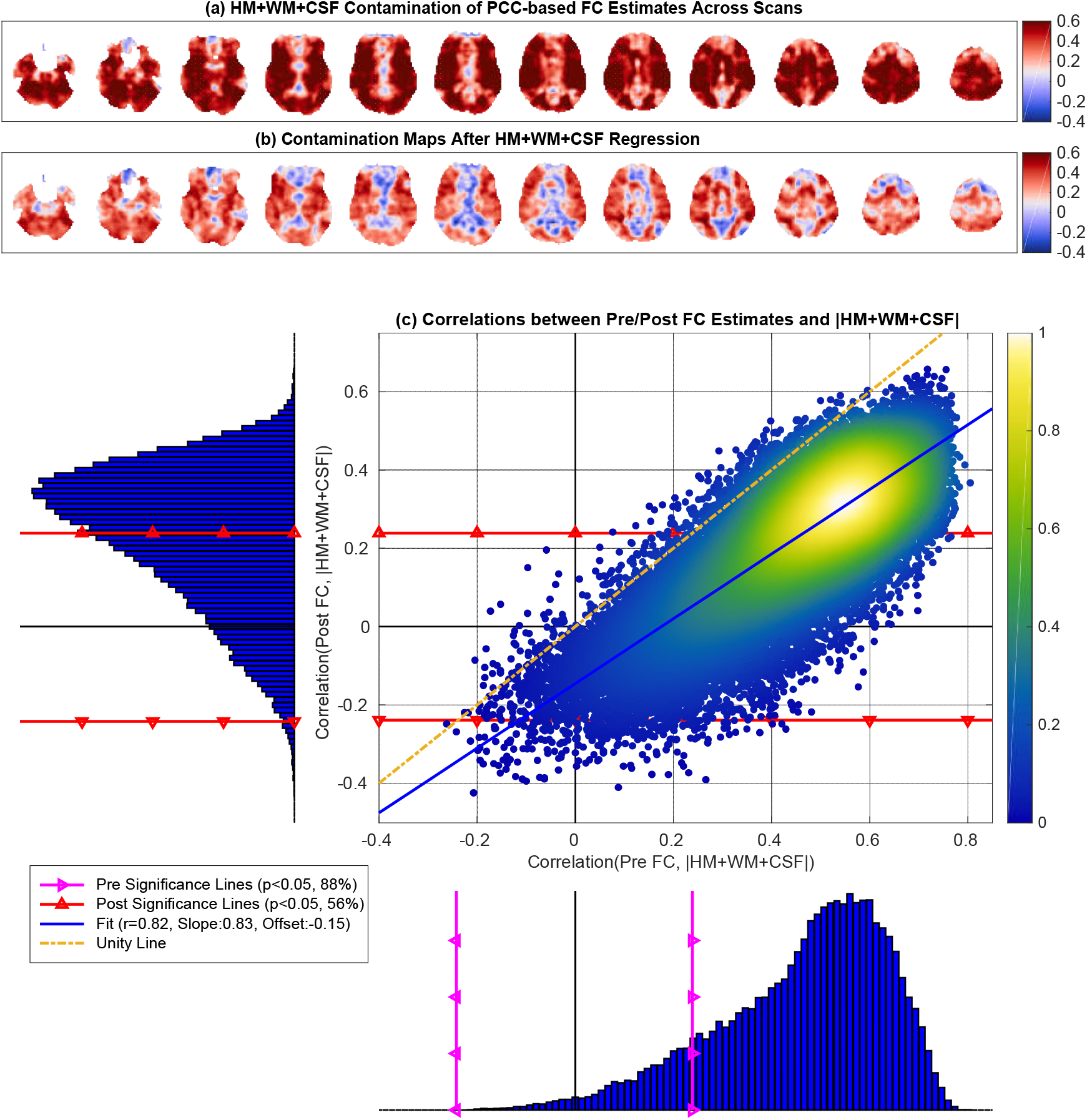
Nuisance contamination maps obtained by correlating the total HM+WM+CSF norm with the FC estimates (a) before and (b) after nuisance regression. In (c) the correlations between the Pre FC estimates and the nuisance norm (x-axis) ranged from = −0.26 to *r* = 0.81 with mean 0.46. The correlations between the Post FC estimates and nuisance norm (y-axis) ranged from = −0.42 to *r* = 0.66 with mean 0.23. There was a strong linear relation between the two correlation distributions (linear fit *r* = 0.82, *p* < 10^−6^). The linear fit between the two correlation distributions was close to the line of unity with a large slope (0.83) and a small negative offset (−0.15).

In Figure 6c the correlations between the Pre FC estimates and nuisance norm ranged from *r* = −0.26 to *r* = 0.81 with mean 0.46. After nuisance regression, the correlation between the Post FC estimates and nuisance norm ranged from *r* = −0.42 to *r* = 0.66 with mean 0.23. We found a strong linear relation between the two correlation distributions (*r* = 0.82, *p* < 10^−3^) with a slight increase in significant anticorrelations residing below the lower red significance line. The linear fit between the two correlation distributions was close to the line of unity with a large slope (0.83) and a small negative offset (−0.15).

As noted in Table 1, 88% of the Pre FC estimates were significantly correlated (*p* < 0.05) with the nuisance norm. The nuisance norm explained an average of 27%±12% (mean+SD) of the variance in those significant correlations. After regression, 56% of the Post FC estimates were still significantly correlated with the nuisance norm, and the nuisance norm explained an average of 13%±6% (mean+SD) of the variance in those Post FC estimates.

In Figure 5b,c we show the ΔFC plots for the WM and CSF regressors. We found that the mean orthogonal nuisance fraction for the WM and CSF regressors were still relatively large with |*n_O_*|^2^/|*n*|^2^ =0.81 and |*n_O_*|^2^/|*n*|^2^ = 0.71, respectively. Due to the theoretical bounds associated with the large orthogonal fractions, the ΔFC points were clustered close to 0 with mean ΔFC values of −0.08 and −0.13 for the WM and CSF regressors, respectively. In Figure 5d we show the ΔFC plot for the HM+WM+CSF regression using the first PC of those regressors. This revealed a mean orthogonal fraction of |*n_O_*|^2^/|*n*|^2^ = 0.73 (similar to individual WM and CSF regressors) with mean ΔFC around −0.13 and ΔFC values bounded below by the theoretical bound. In Supplementary Figure 2, we show the ‘true’ ΔFC values (i.e. not approximated with the 1^st^ PC) were within the theoretical bounds with a mean ΔFC = −0.17, similar to Figure 5d.

To summarize, although HM+WM+CSF regression partially reduced the correlations between the nuisance norm and Pre FC estimates (as compared to HM regression alone), a large fraction of seed-voxel pairs (56%) exhibited significant correlations between the nuisance norm and Post FC estimates. The limited efficacy of nuisance regression reflects the fact that the theoretical bounds on ΔFC estimates in Figure 5d are still very tight since the orthogonal nuisance fraction is large and close to 1.0.

We present the HM+WM+CSF contamination maps and their respective correlation distributions before and after nuisance regression for other seeds (e.g. MOT, AUD, FEF, and IPS) in Supplementary Figures 7 to 10. These were similar to the PCC-based results discussed above. Additionally, Table 1 summarizes the relations between nuisance norms and FC estimates for other seeds.

### 3.3. GS regression

In Figure 7a,b we show GS contamination maps both before and after GSR. In panel (a) we observe strong positive correlations (red regions) between the GS norm and Pre FC estimates. The distribution of these correlations is shown in Figure 8, with values ranging from *r* = −0.22 to *r* = 0.87 with mean 0.57. Across the sample, 96% of these correlations were significant (*p* < 0.05) and 99% were positive with a strong left skew *S* = −0.96. The GS contamination map after GSR is given in Figure 7b. This map shows brain regions consisting of both positive (red regions) and anti-correlations (blue regions) between the GS norms and Post FC estimates.

**Figure 7:**
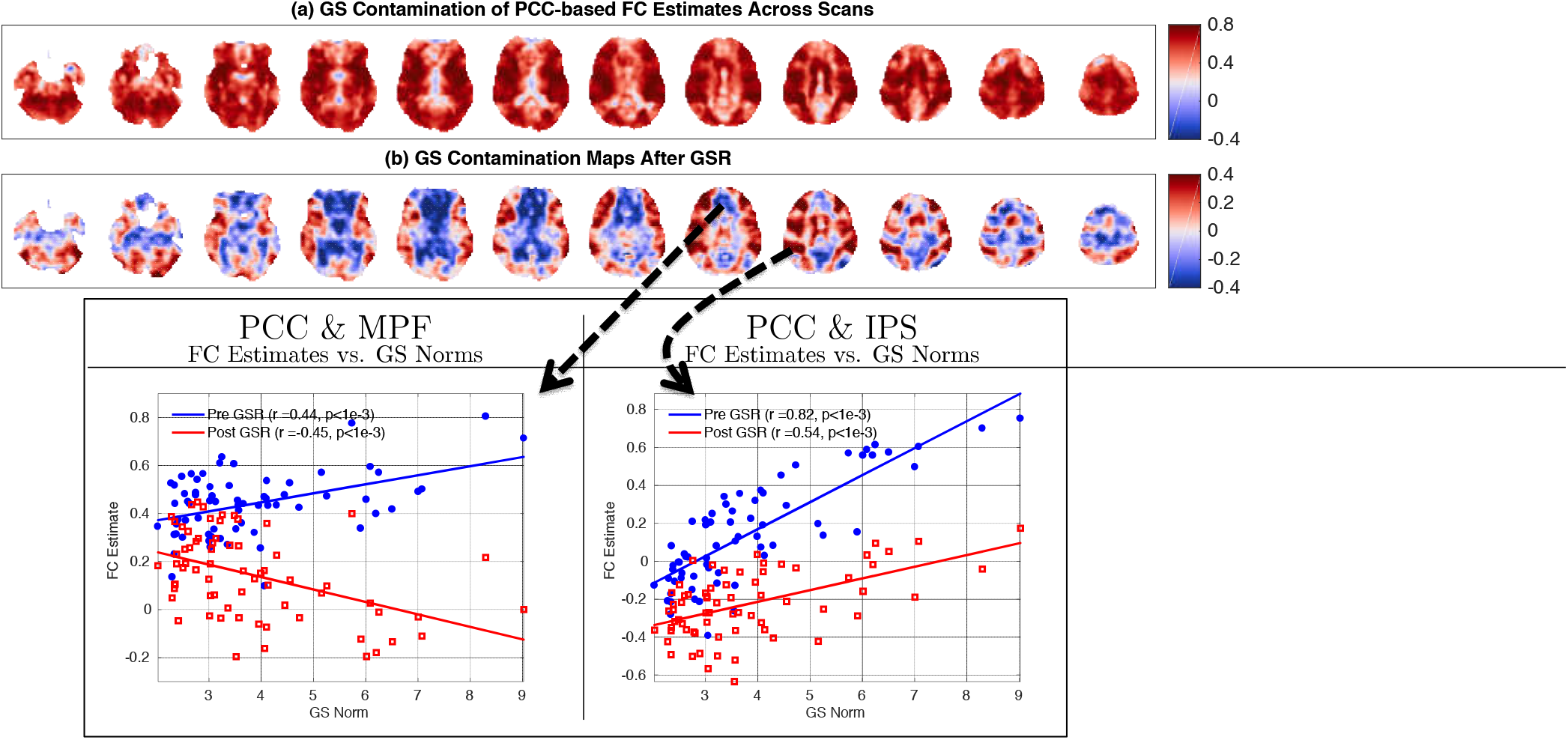
GS contamination maps obtained both (a) before and (b) after GSR. In panel (a) the contamination maps show positive and significant, correlations (99% positive and 96% significant) between Pre FC estimates and GS norm. In (b) the contamination map after GSR involves both positive (49%) and negative (51%) correlations between the GS norms and Post FG estimates. 27% of these correlations were significant (*p* < 0.05). In the seed-pair examples shown at the bottom, the FG estimates between the PGG & MPF across scans are positively correlated with GS norms before GSR (*r* = 0.44) but become ant.i-correlated after GSR (*r* = −0.45). The FG estimates between PGG & MPF seed pair are positively correlated with the GS norm both before and after GSR with correlation values of *r* = 0.82 and *r* = 0.54, respectively.

In Figure 7b we observe that GSR introduces anti-correlations between the GS norm and FC estimates which were not present prior to GSR in panel (a). An example seed pair is provided at the bottom of Figure 7 as a scatter plot. The FC estimates obtained between the PCC&MPF seeds on the left-hand side are positively correlated with the GS norm before GSR and anti-correlated with the GS norm after GSR. As shown along the y-axis of Figure 8 the correlations between the GS norm and Post FC estimates ranged from *r* = −0.61 to *r* = 0.66 centered around a mean 0 with standard deviation 0.21. 49% of these correlations were positive (remaining 51% were negative) and 27% were significant (*p* < 0.05).

**Figure 8:**
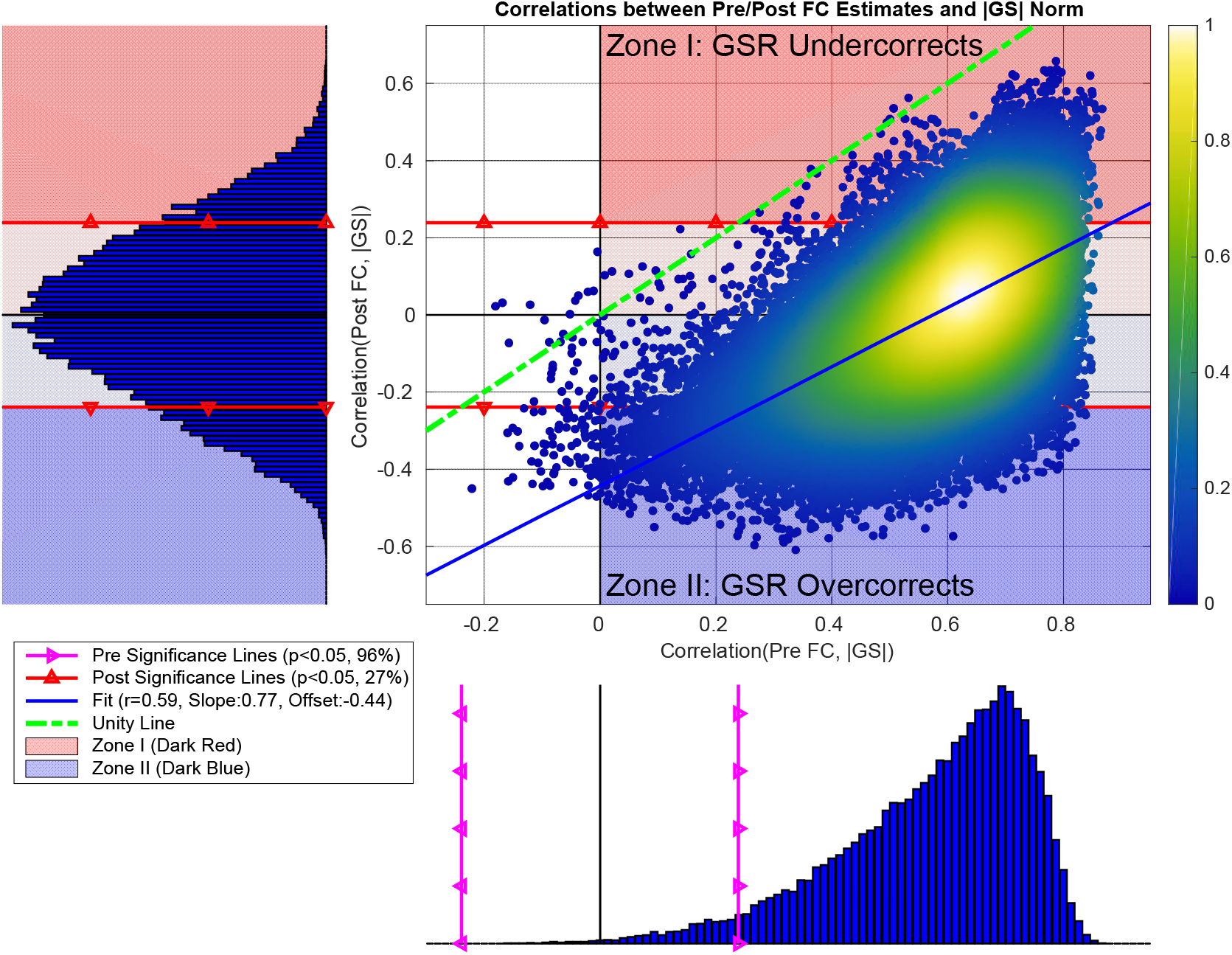
In the scatter plot, we show the correlations between the Post FC estimates and GS norms plotted versus the correlations obtained between the Pre FC estimates and GS norms. We found a significant linear relationship between the two correlation distributions (*r* = 0.59,*p* < 10^−6^). The linear fit (blue line) had a slope of 0.77 and a very large negative offset −0.44. In the bottom histogram, the correlations between the Pre FC estimates and GS norms ranged from *r* = −0.22 to *r* = 0.87 with mean 0.57 and standard deviation 0.16. 99% of the correlations were positive with a strong left skew (Skewness = −0.96) and 96% of the correlations were significant (*p* < 0.05) (correlations residing to the right of magenta line with triangles). In the sideways histogram on the left, the correlations between the Post FC estimates and GS norm ranged from *r* = −0.61 to *r* = 0.66 with mean 0 and standard deviation 0.21, where significance lines (*p* < 0.05) are shown with red lines with triangles. We found 27% of the Post FC estimates to be significantly correlated with HM norms after regression. The labels on the scatter plot (“Zone I” (red zone) and “Zone II” (blue zone)) refer to two main cases where GSR fails to remove significant correlations between the GS norm and Post FC estimates.

In Figure 8 we found a significant linear relationship (*r* = 0.59,*p* < 10^−6^) between the correlations obtained between the Post FC estimates and GS norm versus those correlations obtained before GSR. The linear fit (shown with the blue line) is fairly parallel to the line of unity (green dashed line) with a slope of 0.77 and has a very large negative offset −0.44. The negative offset indicates that GSR produces a strong negative shift in the correlations obtained between the Post FC estimates and GS norms when compared to those correlations obtained before GSR. This results in significant anti-correlations (*p* < 0.05) between the GS norms and Post FC estimates. The significant anti-correlations between the GS norms and Post FC estimates are shown with the points residing below the bottom significance line (red line with triangles) within the dark blue zone in the scatter plot. The points residing above the top significance line in the dark red zone are significant positive correlations remaining after GSR. These correlation values remain significant and positive despite the negative offset.

These results indicate that effects GS norm can largely remain in the Post FC estimates after GSR. GSR can result in “residual” positive correlations between the GS norm and Post FC estimates and can also “introduce” significant anti-correlations between the GS norm and Post FC estimates due to the negative shift in correlation values. To understand why GSR results in both positive and negative correlations between the GS norm and Post FC estimates we start by defining two zones as illustrated in Figure 8 with dark red and dark blue zones labeled by *Zone I* and *Zone II*.

#### 3.3.1. Zone I (GSR undercorrects): GSR is limited in removing GS norm from FC estimates

In Figure 8, red background colors (both light and dark colors) show those correlations between GS norm and FC estimates that are positive both before and after GSR. In this case, “GSR undercorrects” and is unable to fully remove the positive correlations between the GS norm and FC estimates.

We define Zone I (dark red background) to include the significant positive correlations (*p* < 0.05) between GS norm and FC estimates remaining after GSR. In Figure 9a we plot the average FC estimates (across Zone I seed-voxel pairs) versus the GS norm before GSR (blue line and dots) and after GSR (red line and diamonds), where each point corresponds to a single scan. The correlation between the average Pre FC and GS norm is *r* = 0.87. The correlation between the average Post FC and GS norm after GSR is still large *r* = 0.74. This indicates that GSR is unable to fully remove the GS norm effects from the Pre FC estimates.

**Figure 9:**
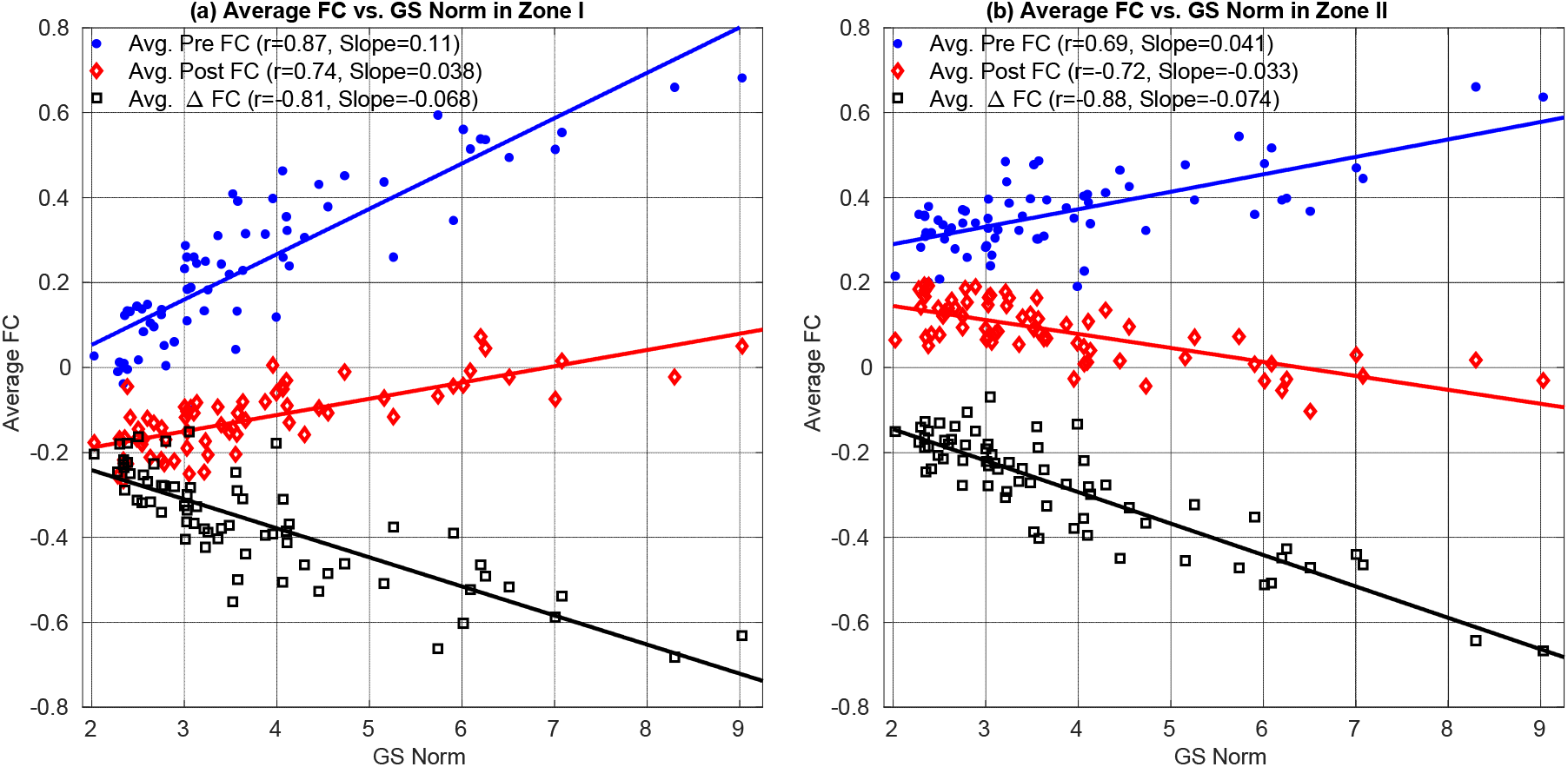
Average FC estimates and average ΔFC versus GS norm in (a) Zone I and (b) Zone II. In both panels the average effect of GSR is equivalent to adding the average ΔFC (shown with black color) to the average Pre FC estimates (blue color). In Zone 1, the resulting average Post FC estimates (red color) are still positively correlated with the GS norm *r* = 0.74. This is because the slope for average ΔFC (−0.068) has a smaller magnitude than the slope for average FC estimates (+0.11). Thus, the slope for average Post FC estimates remains positive 0.038 ≈ +0.11 − 0.068. In panel (b) the average Pre FC estimates for Zone II are significantly larger than those in Zone I in panel (a) *p* < 10^−3^ (paired two-tailed t-test) with a smaller standard deviation (0.09 in Zone II as compared to the 0.19 in Zone I). Thus, the slope for average Pre FC estimates in Zone II has smaller magnitude +0.041 as compared to Zone I. The slope (−0.074) of average ΔFC in (b) is similar to the slope (−0.068) of average ΔFC in (a). Since the average Post FC estimates are simply the sum of average Pre FC points and average ΔFC points, the average Pre FC estimates are dominated by the larger ΔFC effect and the relation between the average Post FC estimates and GS norm exhibits a negative correlation (*r* = −0.72, *p* < 10^−3^).

The effect of GSR is fully characterized by the average ΔFC values (black squares) shown in Figure 9a. The average ΔFC is anti-correlated with the GS norm (*r* = −0.81) where the slope of the linear fit (black line) is −0.068. The slope (0.11) of the linear fit (blue line) between the Pre FC estimates and GS norm has a greater magnitude than the slope (−0.068) for ΔFC. Since Post FC = Pre FC + ΔFC, this difference in the slope magnitudes results in a positive slope for the relation between Post FC estimates and GS norm (linear fit shown with red line).

In Figure 10a we plot the ΔFC values versus orthogonal nuisance fraction for Zone I, where each point represents the values obtained for a single seed-voxel pair in a given single scan. The ΔFC values are strongly related to the orthogonal nuisance fraction with *r* = 0.93 (*p* < 10^−3^), where the linear fit shown with the green line is almost tangent to the lower theoretical bound. We observe that ΔFC values are clustered around the lower theoretical bound with mean value of −0.37 with a relatively smaller standard deviation −0.17. This is consistent with the average ΔFC values in Figure 9a. The bound plot reveals that the effects of GSR on the FC estimates in Zone I are bounded below by the theoretical curve such that the magnitudes of ΔFC cannot exceed the lower bound. As a result, the magnitude of ΔFC in Figure 9a cannot increase as rapidly as the Pre FC estimates.

**Figure 10:**
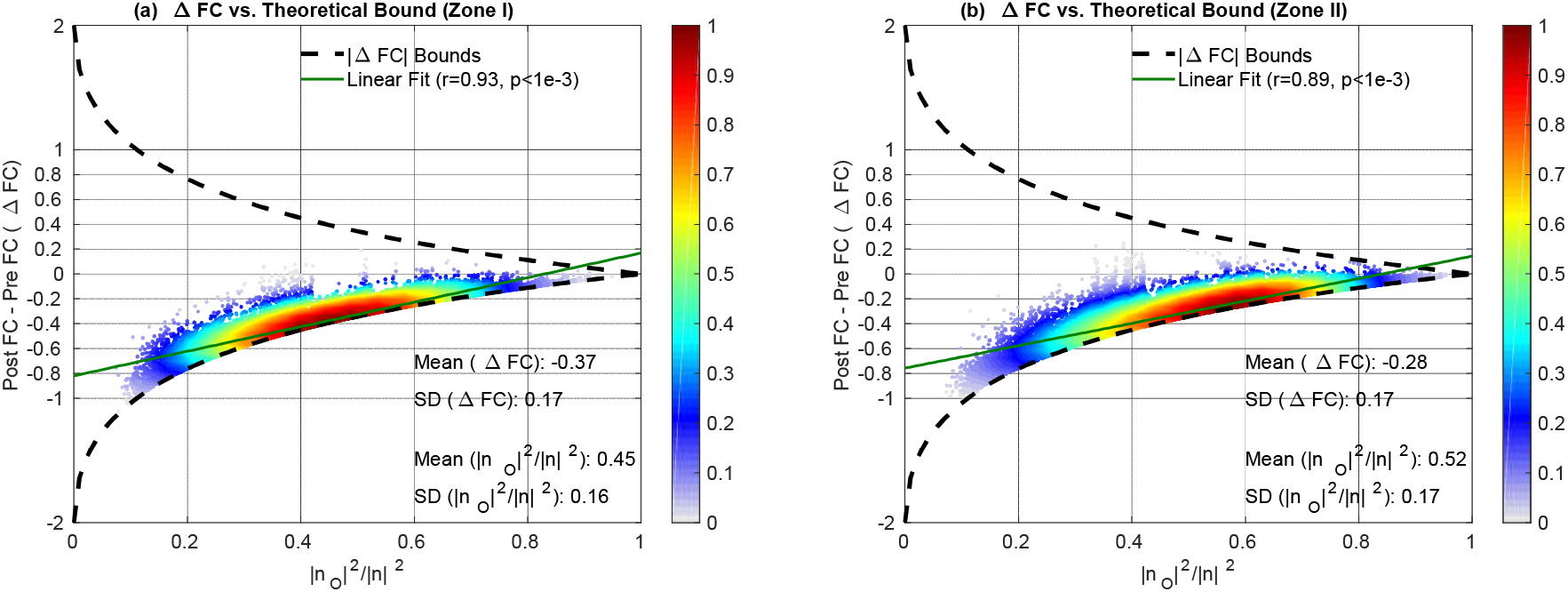
ΔFC versus orthogonal nuisance fraction 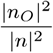 for GS regression in (a) Zone I and (b) Zone II. In both zones, ΔFC values are clustered close to the lower theoretical bound with mean ΔFC −0.37 for Zone I and −0.28 for Zone II with similar mean orthogonal nuisance fractions of 0.45 and 0.52. Moreover, the standard deviation for ΔFC is 0.17 which is much smaller than the mean ΔFC values. The linear fits shown with solid green lines show that ΔFC values are strongly correlated (*p* < 10^−3^) with the orthogonal nuisance fractions with *r* = 0.93 and *r* = 0.89 for Zone I and Zone II, respectively. Both linear fits are extremely close to the lower theoretical bound.

#### 3.3.2. Zone II (GSR overcorrects): GSR introduces anti-correlation between GS norm and FC estimates

The light and dark blue backgrounds in Figure 8 show those correlations between GS norm and FC estimates that are **positive** before GSR and are **negative** after GSR. In this case, “GSR overcorrects” by first removing the positive correlations between GS norm and Pre FC estimates and then introducing anti-correlations between the GS norms and Post FC estimates. The brain regions which are red in Figure 7a but are blue in Figure 7b correspond to this case. These regions broadly include the default mode network, medial-prefrontal cortex, and thalamic regions, when using the PCC as the seed signal.

In Figure 8 we define Zone II (dark blue background) to include **significant** negative correlations (*p* < 0.05) between GS norm and FC estimates after GSR. In Figure 9b we plot the average FC estimates versus the GS norm for Zone II. The slope (0.041) of the relation between Pre FC estimates and GS norm in Zone II in Figure 9b is weaker in magnitude as compared to the slope (−0.074) of the relation between the ΔFC and GS norm. Since Post FC = Pre FC + ΔFC, a stronger negative relation between ΔFC and GS norm dominates over the weaker relation between Pre FC estimate and GS norm and the average Post FC estimates become significantly anti-correlated with the GS norm (*r* = −0.72, *p* < 10^−3^).

The average Pre FC estimates in Zone II in Figure 8b are significantly greater than those in Zone I in panel (a) *p* < 10^−3^ (paired two-tailed t-test). This means that BOLD signals residing in Zone II exhibit greater intrinsic similarity (on average) to the seed signal as compared to those in Zone I. To verify this, in Figure 11a we plot the correlations between the seed signal (PCC) and the average BOLD signal in Zone II versus the correlations between the seed signal and the average BOLD signal in Zone I. The average BOLD signal in Zone II was significantly (*p* < 10^−3^ paired two-tailed t-test) more correlated with the seed signal as compared to Zone I.

**Figure 11:**
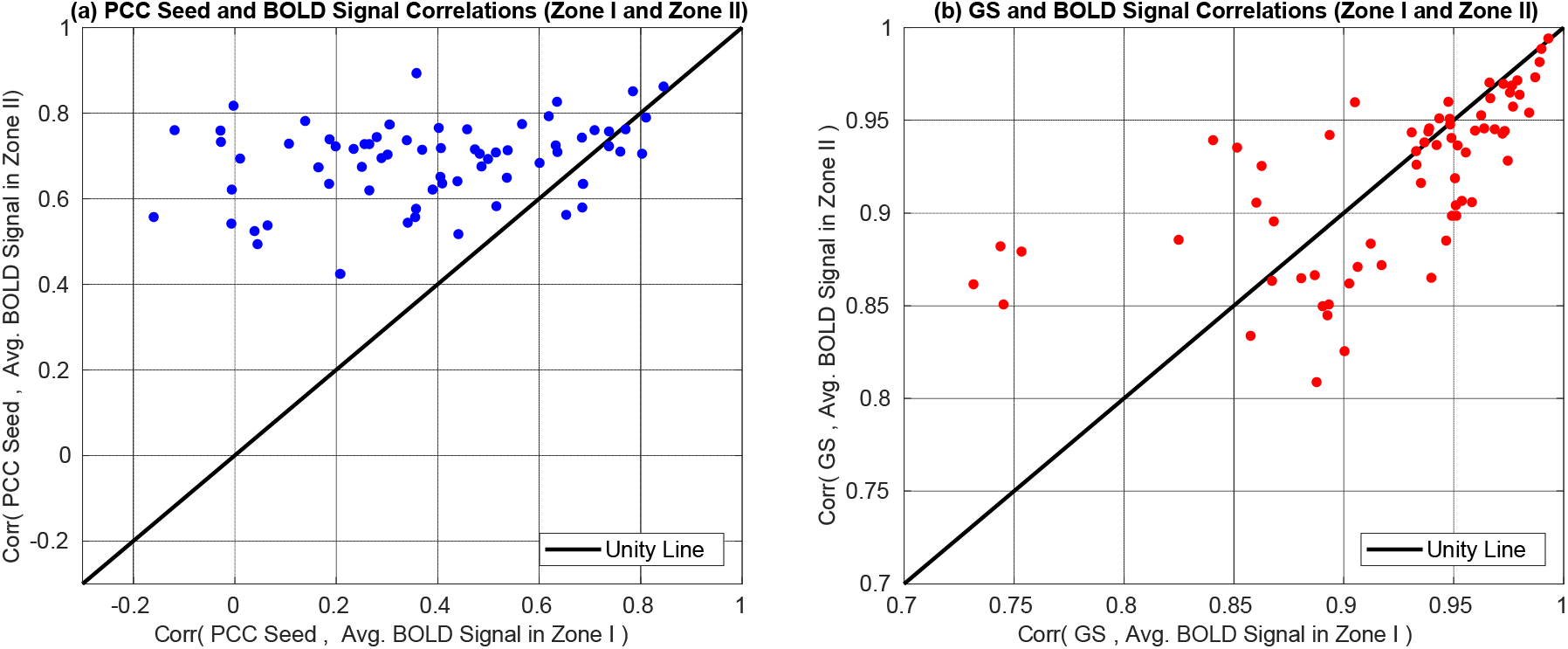
In (a) we show the correlations obtained between the PCC seed and the average BOLD signal in Zone II versus the correlations obtained between the PCC seed and the average BOLD signal in Zone I. The average BOLD signal in Zone II was significantly (*p* < 10^−3^ paired two-tailed t-test) more correlated with the PCC seed signal as compared to Zone I. This means that brain regions that are more similar to the seed signal are more likely to have the Zone II limitation of GSR. In (b) we show the correlations between the GS the average BOLD signal in Zone II versus the correlations between the GS and the average BOLD signal in Zone I. We found no significant difference (*p* = 0.45, paired two-tailed t-test) between Zone I and Zone II correlations. This indicates that on average, the differences in Zone I and Zone II effects are not dependent on the relation between the GS and the underlying BOLD time courses in those zones.

In Figure 11b we plot the correlations between the GS and the average BOLD signal in Zone II versus the correlations between the GS time course and the average BOLD signal in Zone I. We found no significant difference (*p* = 0.45, paired two-tailed t-test) between those correlation values for Zone I and Zone II. This means that the GS regressor exhibits a similar range of correlations to the raw time courses in both zones. This is consistent with the fact that the average ΔFC values in the two zones (black squares in Figure 9) were highly correlated across scans with each other (*r* = 0.94). In addition, the relation between the per-voxel ΔFC values and the orthogonal nuisance fractions were similar in the two zones as shown by the linear fits in Figure 10 with *r* = 0.93 for Zone I and *r* = 0.89 for Zone II.

In summary, the difference in Zone I and Zone II behavior reflects two effects. First, GSR results in a similar range of ΔFC values for both zones as shown in Figure 10a,b. This was largely because the GS time courses were similarly related to the average BOLD signals in Zone I and Zone II (Figure 11b) and thus the orthogonal nuisance fraction fractions between the GS and seed-voxel pairs were similar in Figure 10 for both zones. Second, FC estimates in Zone II were significantly greater as compared to those in Zone I and exhibited a weaker dependency on the GS norm (slope 0.041, *r* = 0.69) in Figure 9b as compared to the Zone I (slope 0.11, *r* = 0.87).

Figure 11a shows that brain regions that are intrinsically similar to the seed signal are more likely to belong to Zone II. The specific brain regions that exhibit either Zone I limitation (residual positive correlation between the GS norm and Post FC estimates) or Zone II limitation (an introduced negative correlation between the GS norm and Post FC estimates) vary with the seed signal used. These regions can be determined by looking at the GS contamination maps after GSR for other seeds as provided in the Supplementary Figures 11 to 14. Table 1 summarizes the relation between the GS norm and FC estimates.

As a final supplementary experiment, we performed multiple HM+WM+CSF+GS regression. We provide the nuisance contamination maps and correlations in Supplementary Figure 15 using the PCC seed. Both the contamination maps, correlation distributions, and significance results in Table 1 for the multiple regression were very similar to that of GSR.

## 4. Discussion

We have shown that inter-scan variations in FC estimates can be strongly and significantly correlated with the geometric norms of various nuisance measurements. We found that the relationship between the FC estimates and nuisance norms can persist even after performing multiple nuisance regression. We used the mathematical framework developed in (Nalci et al., 2019) to describe the limitations of nuisance regression with regards to static FC measures and demonstrated that the empirical results were in agreement with the theoretically predicted bounds.

For non-GS regressors (i.e. HM and HM+WM+CSF), we found nuisance regression to be largely ineffective. This was because non-GS regressors were largely orthogonal to the measured BOLD data, as shown in Figure 5, with mean orthogonal nuisance fractions ranging from 0.70 to 0.99. The large orthogonality imposed tight bounds on the difference between the FC estimates obtained before and after nuisance regression, which in turn limited the ability of regression to reduce the strength of the correlation between the nuisance norms and the FC estimates. This limitation for non-GS regression applied to static FC measures is largely consistent with our previous findings for dynamic FC measures (Nalci et al., 2019).

We introduced *nuisance contamination maps* to visualize the effects of nuisance norms on the FC estimates both before and after nuisance regression. These maps serve as a useful tool in analyzing the spatial location and extent of nuisance contamination present in seed-based FC estimates across scans. In Figures 3 and 6, the FC estimates obtained between the PCC seed and BOLD signals from other brain regions exhibited strong correlations with the HM and HM+WM+CSF norms both before and after nuisance regression. As summarized in Table 1, depending on the seed signal, 40-100% of the seed-voxel pairs in the brain exhibited significant correlations with respective nuisance norms prior to nuisance regression. After nuisance regression, FC estimates from 43-82% of the seed-voxel pairs still exhibited significant correlations with nuisance norms.

### 4.1. GSR Specific Findings

We found that GSR removed a large portion of the GS norm fluctuations from the FC estimates across scans. However, depending on the seed signal, a considerable portion (20-30%) of the FC estimates still remained significantly correlated with the GS norm. In addition, GSR introduced a negative correlation between the FC estimates and GS norm in some voxels.

To better understand the behavior of GSR, we divided the brain into two spatially non-overlapping regions, corresponding to Zones I and II. The changes in FC (ΔFC) caused by GSR showed a similar negative dependence on the GS norm in both zones. Although the Pre FC values in both zones showed a positive dependence on the GS norm, the slope was smaller for the Zone II voxels, reflecting a weaker dependence. Due to this difference in slopes, significant negative correlations between the GS norm and inter-scan FC estimates were observed after GSR in Zone II, whereas the GS norm remained significantly positively correlated with inter-scan FC estimates in Zone I.

The GS contamination maps obtained with the PCC seed in Figure 7b resembled PCC-based FC maps obtained after GSR (Fox et al., 2005; Murphy and Fox, 2016). It is known that GSR increases the spatial extent and strength of anti-correlations present in resting-state FC maps, especially between the default-mode (DMN) and task-positive networks (TPN) (Fox et al., 2005; Saad et al., 2012; Fox et al., 2009; Murphy et al., 2009; Murphy and Fox, 2016). Indeed a great deal of the controversy concerning the use of GSR has centered on whether the observed anti-correlations are real or artifactual (Murphy and Fox, 2016; Nalci et al., 2017). Despite the visual similarity of the two types of maps, it is important to stress that they depict two fundamentally different quantities, with the contamination maps showing the relation between the GS norm and FC estimates across scans.

To better understand the similarity in the appearance of the two types of maps, we first note that the weaker dependence of the Zone II Pre FC values on the GS norm was associated with the presence of significantly higher Pre FC values in this zone as compared to Zone I. This association suggests that the FC measures for voxels that are more strongly correlated with the seed signal exhibit a weaker sensitivity to variations in the norm of the GS as compared to voxels with a weaker correlation. For the PCC seed, the TPN region resided in Zone I and the DMN region resided in Zone II, consistent with a stronger correlation with the PCC for voxels in the DMN. In both zones, GSR produced a similarly sized negative offset (mean of −0.37) on the Pre FC estimates. The observation of a negative offset is consistent with prior findings (Murphy et al., 2009). As a result of this negative offset and the underlying differences in the Pre FC values, the mean FC estimates in Zone I in Figure 9a became negative after GSR, while the FC estimates in Figure 9b remained largely positive. To summarize, Zone I voxels exhibit both a positive dependence on GS norm and negative FC values after GSR, whereas Zone II voxels exhibit both a negative dependence on GS norm and positive FC values after GSR. These associations give rise to the similarity in the maps.

The current findings add additional factors to consider in the ongoing controversy regarding the use of GSR (Murphy and Fox, 2016; Nalci et al., 2017; Liu et al., 2017; Glasser et al., 2018). Prior to GSR a high percentage (>96%) of the FC estimates were significantly correlated with the GS norm, with the GS norm accounting for 40 ± 12 percent of the variance in the whole-brain FC estimates across scans and different seeds. This means that differences in GS norms can largely drive variations in FC metrics across scans when GSR is not performed. As discussed below, this dependence may partly reflect variations in vigilance across scans. On the other hand, when GSR is performed, the strength of the relation between the FC estimates and the GS norms is reduced, but there is a risk of introducing negative correlations between the GS norm and inter-scan FC estimates. Further work is needed to assess the impact of these negative correlations on the interpretation of FC studies.

### 4.2. Studies Investigating FC Across Scans

A multitude of fMRI studies have investigated the differences in FC measures between disease populations and healthy controls. Examples of such studies include investigations of Alzheimer’s disease (Greicius et al., 2004; Yang et al., 2014; Wang et al., 2007), Parkinson’s disease (Baudrexel et al., 2011), depression (Greicius et al., 2007), schizoprenia (Liu et al., 2008), dementia (Rombouts et al., 2009), and amyotrophic lateral sclerosis (ALS) (Mohammadi et al., 2009; Agosta et al., 2013). While these studies all used some type of nuisance regression to remove the potential effects of nuisance terms on their analyses, our results strongly indicate that the results may still have exhibited a relation between the FC estimates and the nuisance norms.

As an example, in (Baudrexel et al., 2011) the authors observed an increased subthalamic nucleus-motor cortex FC in Parkinson’s disease as compared to healthy controls. They compared HM measures across the Parkinson’s and healthy control groups but did not find a significant difference. However, they did not directly consider either the relation between HM measures and FC estimates or potential group differences in the variation of HM parameters across each group. We have shown that HM norms can be strongly correlated with the FC estimates across scans both before and after regression. Based on our findings, it is possible that inter-group differences in the variability of HM measures could have affected the finding of FC differences. We believe that any future study that compares FC estimates across subject groups should also consider the potential link between FC estimates and nuisance norms.

### 4.3. Limitations of Linear Regression

fMRI studies typically rely on linear regression to remove nuisance effects from BOLD data by either using direct nuisance measurements (Bright et al., 2017; Power et al., 2012; Glover et al., 2000; Power et al., 2015; Chang et al., 2009; Chang and Glover, 2009) or using a set of nuisance regressors derived with data-driven methods (Behzadi et al., 2007; Pruim et al., 2015; Glasser et al., 2018). The performance of these methods is usually characterized by their effects on raw BOLD signals on a per-scan basis (e.g. the percentage of variance removed). The ability of these techniques to reduce the relation between FC estimates and nuisance norms has not been as fully explored (Nalci et al., 2019). As shown in both (Nalci et al., 2019) and the present work, the ability of linear regression methods to reduce the strength of the relation between FC estimates and nuisance norms can be quite limited even when the regressor exhibits only a moderate degree of orthogonality to the BOLD data. This limitation holds regardless of whether a regressor is obtained using data-driven methods or measured directly. Further work is needed to develop nuisance mitigation methods that can go beyond the limitations of linear regression approaches.

### 4.4. Nuisance Norm Regression

The present study shows the limited efficacy of nuisance regression in reducing the relation between nuisance norms and FC estimates across scans. Given this limitation, a reasonable approach is to perform nuisance norm regression (NNR) on the inter-scan FC estimates. This approach was originally proposed in Nalci et al. (2019) for cleaning sliding-window dynamic FC estimates and involves projecting out the nuisance norms from inter-scan FC estimates. Though this method would ensure orthogonality between the nuisance norms and inter-scan FC estimates, it might not be a suitable approach for cleaning static FC estimates.

One potential issue with NNR for static FC estimates is the presence of leverage effects (Hoaglin and Welsch, 1978; Draper and Smith, 2014). In computing the regression fit coefficient between the nuisance norms and inter-scan FC estimates, scans with larger nuisance norms will have greater leverage on the regression equation. This means that a scan with a large nuisance norm value can have a relatively greater influence on the fit coefficient. Therefore, although NNR can eliminate the correlation between FC estimates and nuisance norms across scans, there may be potential issues that require further study. Future work and new approaches are of interest to understand how to best minimize nuisance norm effects in inter-scan FC estimates.

### 4.5. Vigilance Effects

There is growing evidence that changes in vigilance are responsible for a considerable portion of the resting-state fMRI signal (Wong et al., 2012, 2013, 2015; Liu et al., 2017; Falahpour et al., 2018). Vigilance fluctuations can account for approximately 10-20% of the variance in the whole-brain GS, while differences in mean vigilance levels can account for 25% of the variance in the GS norm across a sample and 64-81% of the differences in GS norms observed between conditions (Liu et al., 2017). In addition, Falahpour et al. (2018) demonstrated large variations in the vigilance norm (i.e. standard deviation of vigilance fluctuations within a scan) across scans. Our results in Table 1 reveal that inter-scan FC estimates are strongly related to fluctuations in the GS norm across scans. Given the reported relations between the GS and vigilance measures, it is likely that variations in static FC estimates will exhibit a dependence on variations in vigilance. Further work to study the relation between vigilance norms and FC estimates will be helpful.

## 5. Conclusion

We have provided a detailed examination of nuisance effects and the efficacy of nuisance regression on the variability of FC estimates across scans. We have shown that the norms of various nuisance terms can be strongly and significantly correlated with variations in inter-scan FC estimates both before and after nuisance regression. We found that non-GS regressions (HM, HM+WM+CSF) were largely ineffective in reducing the correlations between nuisance norms and FC estimates. We showed that though GSR removed a large fraction of the GS norm fluctuations from the FC estimates, a considerable portion (20 – 30%) of the FC estimates still remained significantly correlated with the GS norm. In addition, significant negative correlations between the GS norm and FC estimates were introduced by the process.

This work stresses an important issue in the interpretation of FC measures. Most FC studies implicitly assume that the effects of nuisance terms on FC estimates are minimized by nuisance regression in the pre-processing stage. Our findings strongly suggest that the interpretation of FC measures should also consider correlations with the nuisance norms that can persist even after nuisance regression. If the relationship between FC estimates and nuisance norms is not considered, differences in FC estimates may be incorrectly interpreted as meaningful effects, when in fact they may be largely due to differences in nuisance activity. This is especially true in studies comparing FC measures between disease groups or treatment conditions.

As we have shown, linear regression-based nuisance removal approaches exhibit theoretical limitations with regards to their ability to reduce the correlations between FC estimates and nuisance norms. Future FC studies will greatly benefit from the development of nuisance removal approaches that can address the limitations highlighted in this work.

## Acknowledgments

This work was partially supported by NIH grant R21MH112155 and a UC San Diego Frontiers of Innovation Scholars Program (FISP) Project Fellowship.

## 6. Supplementary Material

**Supplementary Figure 1:**
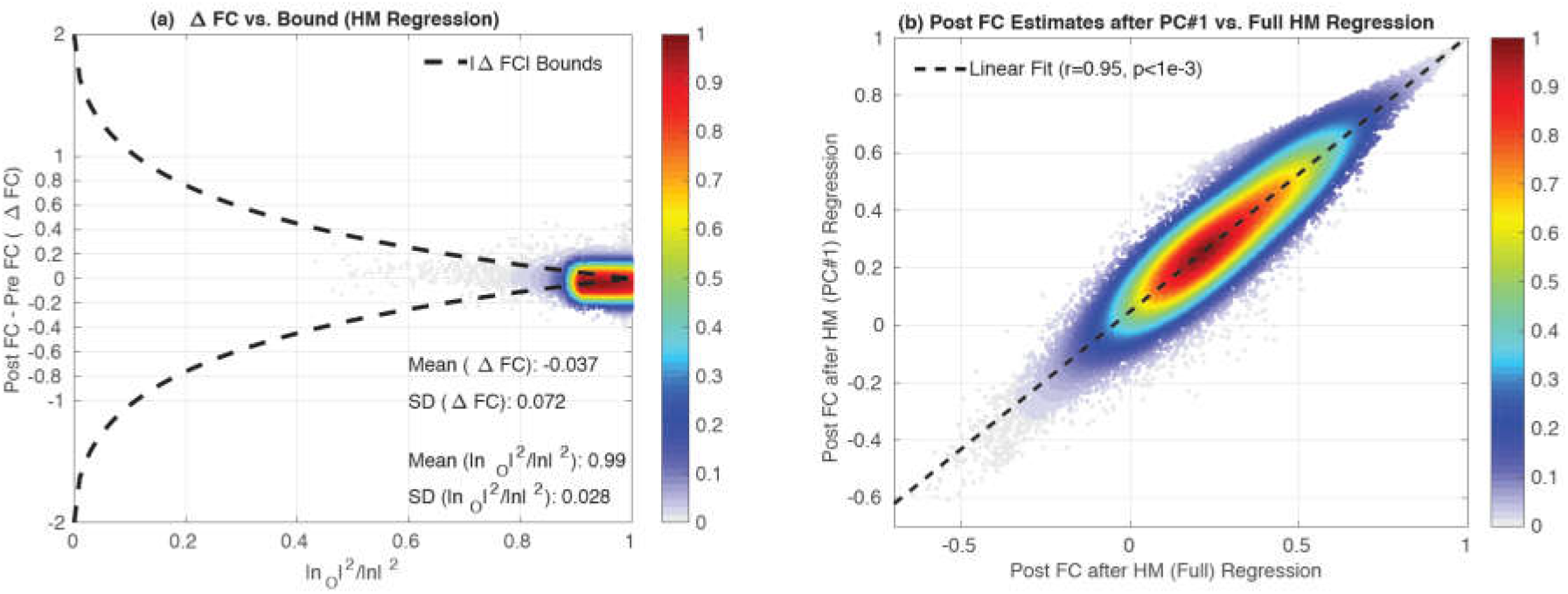
(a) Empirical ΔFC values versus orthogonal nuisance fraction |*n_O_*|^2^/|*n*|^2^ for the 6 HM regressors used in Figure 5a. For this figure full multiple HM regression was performed instead of using the 1^st^ principal component (PC) of the 6 HM regressors to obtain the ΔFC values, as was performed in the main text. The empirical ΔFC values are clustered around a mean value of −0.04 lie within a fairly narrow neighborhood around the range delineated by theoretical bounds. They do not strictly lie within the bounds because the theoretical framework is currently limited to the case of a single regressor and serves only as a rough guide for multiple regressors. (b) Post FC estimates obtained after regressing out the 1^st^ PC of HM measurements and estimates obtained after regressing out all 6 HM regressors were strongly correlated with *r* = 0.95.

**Supplementary Figure 2:**
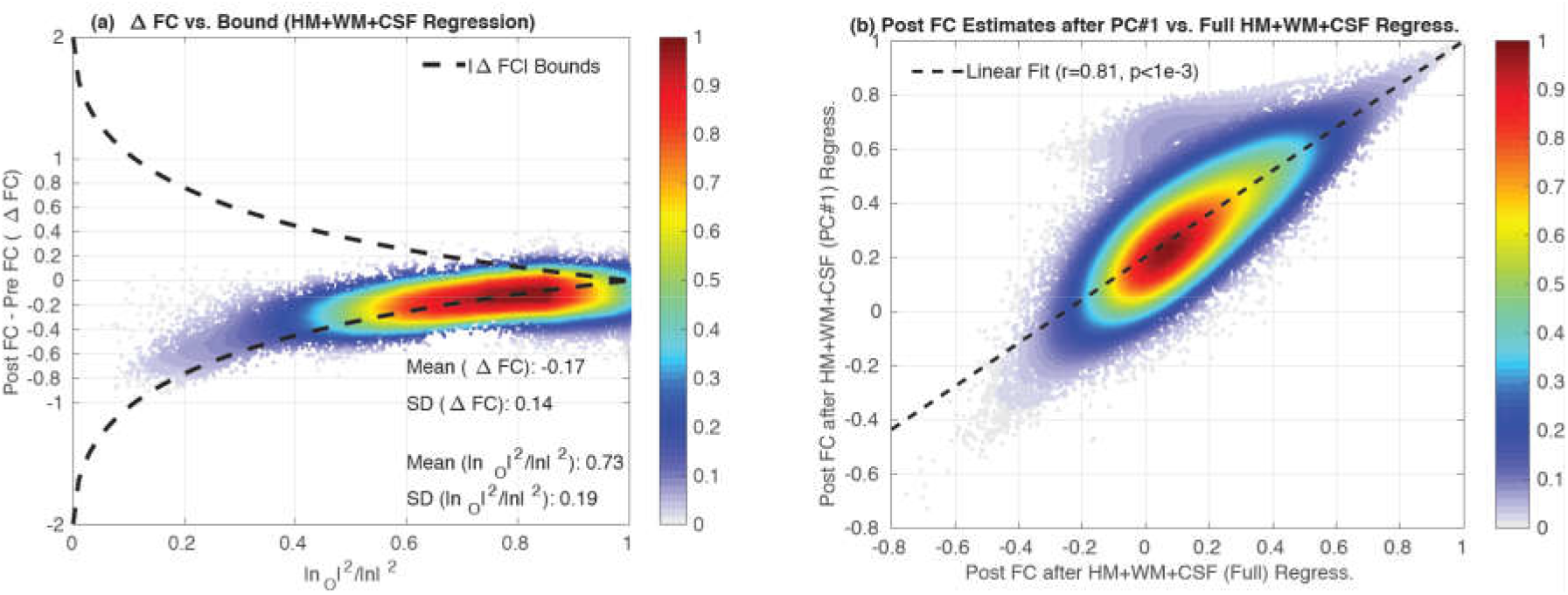
(a) Empirical ΔFC values versus the orthogonal nuisance fraction |*n_O_*|^2^/|*n*|^2^ for the HM+WM+CSF regressors used in Figure 5d. For this example multiple regression was performed instead of using the 1^st^ principal component (PC) of all the regressors to obtain the ΔFC values, as was performed in the main text. The empirical ΔFC values are clustered around a mean value of −0.17 and lie within a fairly narrow neighborhood around the range delineated by theoretical bounds. They do not strictly lie within the bounds because the theoretical framework is currently limited to the case of a single regressor and serves only as a rough guide for multiple regressors. (b) Post FC estimates obtained after regressing out the 1^st^ PC of all HM+WM+CSF regressors and after performing multiple HM+WM+CSF regression were strongly correlated with *r* = 0.81.

**Supplementary Figure 3:**
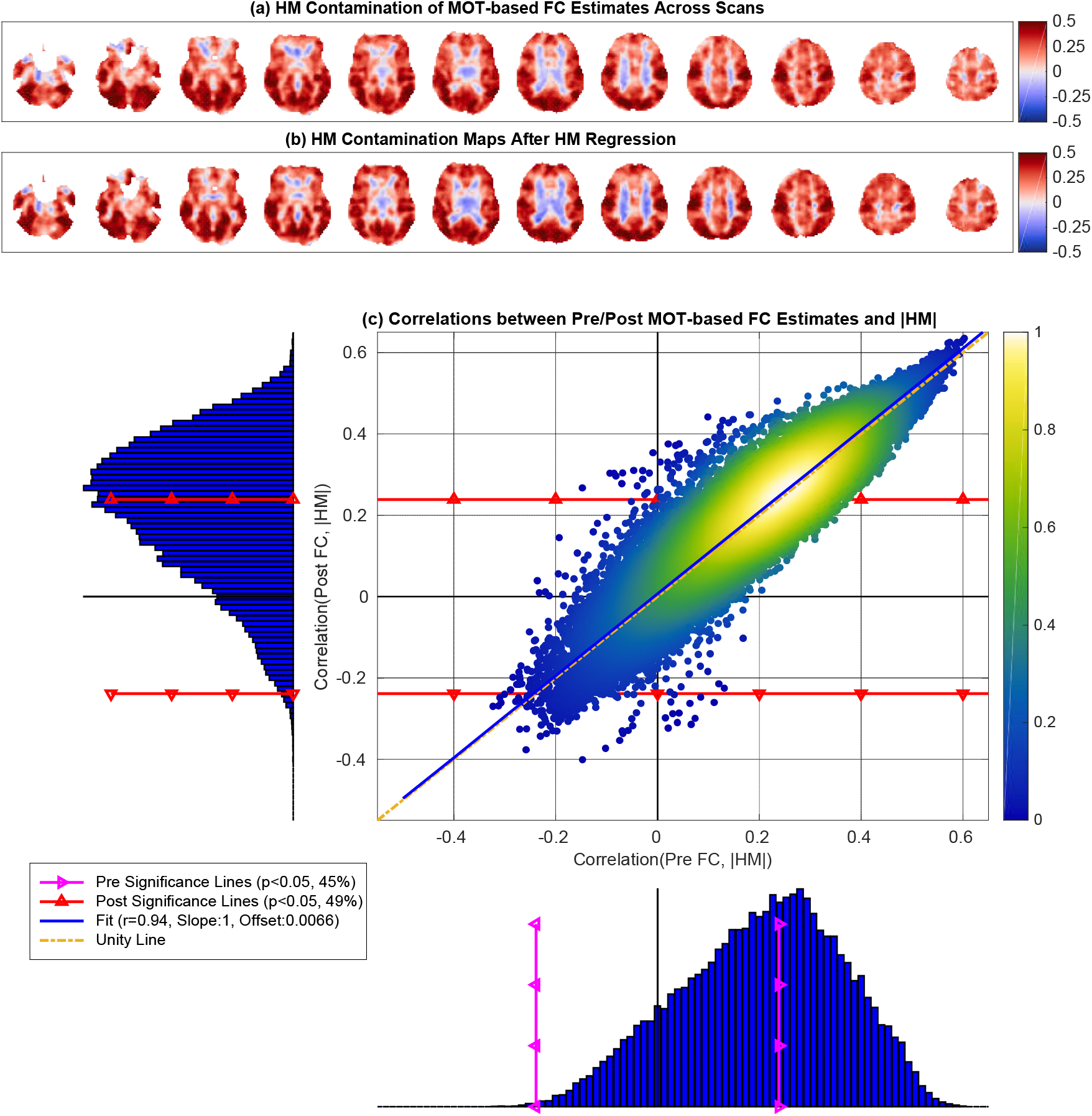
MOT-based HM contamination maps (a) before and (b) after HM regression. These maps were fairly similar to each other (cosine similarity *S* = 0.98) and show widespread correlations between the HM norm and FC estimates across scans. The scatter plot in (c) shows the correlations between the Post FC estimates and HM norm after regression versus the correlations obtained between Pre FC estimates and HM norm. These correlation distributions were significantly related (*r* = 0.94, *p* < 10^−3^) to each other. The linear fit (blue line, Slope= 1 and Offset= 0.007) between the two correlation distributions was nearly identical to the line of unity (dashed yellow line). In the bottom histogram, the correlations between the Pre FC estimates and HM norms ranged from *r* = −0.32 to *r* = 0.60 with mean 0.2, and 45% of these correlations were significant (*p* < 0.05). In the sideways histogram on the left, the correlations between the Post FC estimates and HM norms ranged from *r* = −0.40 to *r* = 0.64 with mean 0.21. The post regression significance lines are shown with red lines with triangles. 49% of the Post FC estimates were significantly correlated with HM norms after regression.

**Supplementary Figure 4:**
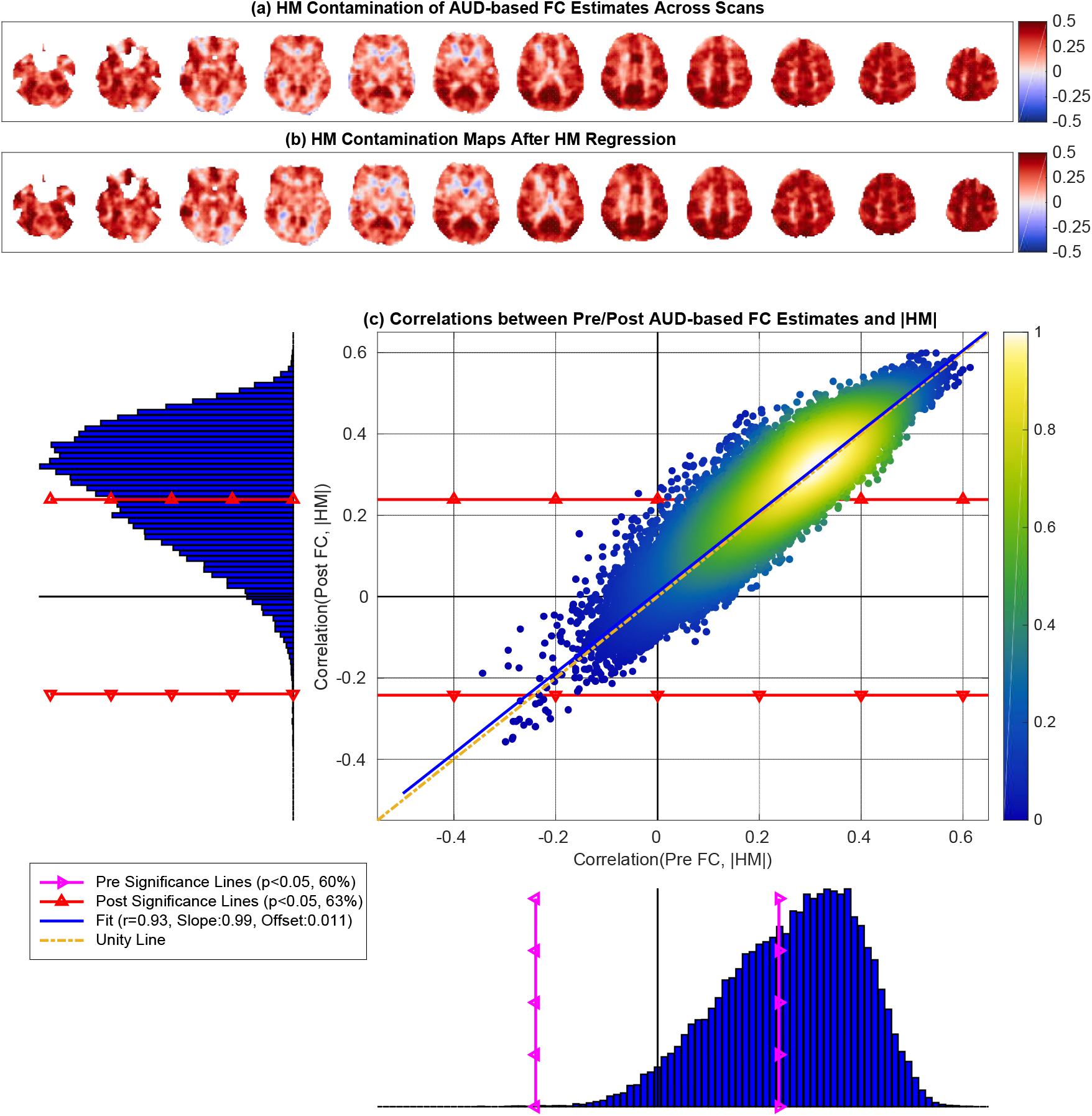
AUD-based HM contamination maps (a) before and (b) after HM regression. These maps were fairly similar to each other (cosine similarity *S* = 0.99) and show widespread correlations between the HM norm and FC estimates across scans. The scatter plot in (c) shows the correlations between the Post FC estimates and HM norm after regression versus the correlations obtained between Pre FC estimates and HM norm. These correlation distributions were significantly related (*r* = 0.93, *p* < 10^−3^) to each other. The linear fit (blue line, Slope= 0.99 and Offset= 0.01) between the two correlation distributions was nearly identical to the line of unity (dashed yellow line). In the bottom histogram, the correlations between the Pre FC estimates and HM norms ranged from *r* = −0.34 to *r* = 0.61 with mean 0.26, and 60% of these correlations were significant (*p* < 0.05). In the sideways histogram on the left, the correlations between the Post FC estimates and HM norms ranged from *r* = −0.36 to *r* = 0.60 with mean 0.27. The post regression significance lines are shown with red lines with triangles. 63% of the Post FC estimates were significantly correlated with HM norms after regression.

**Supplementary Figure 5:**
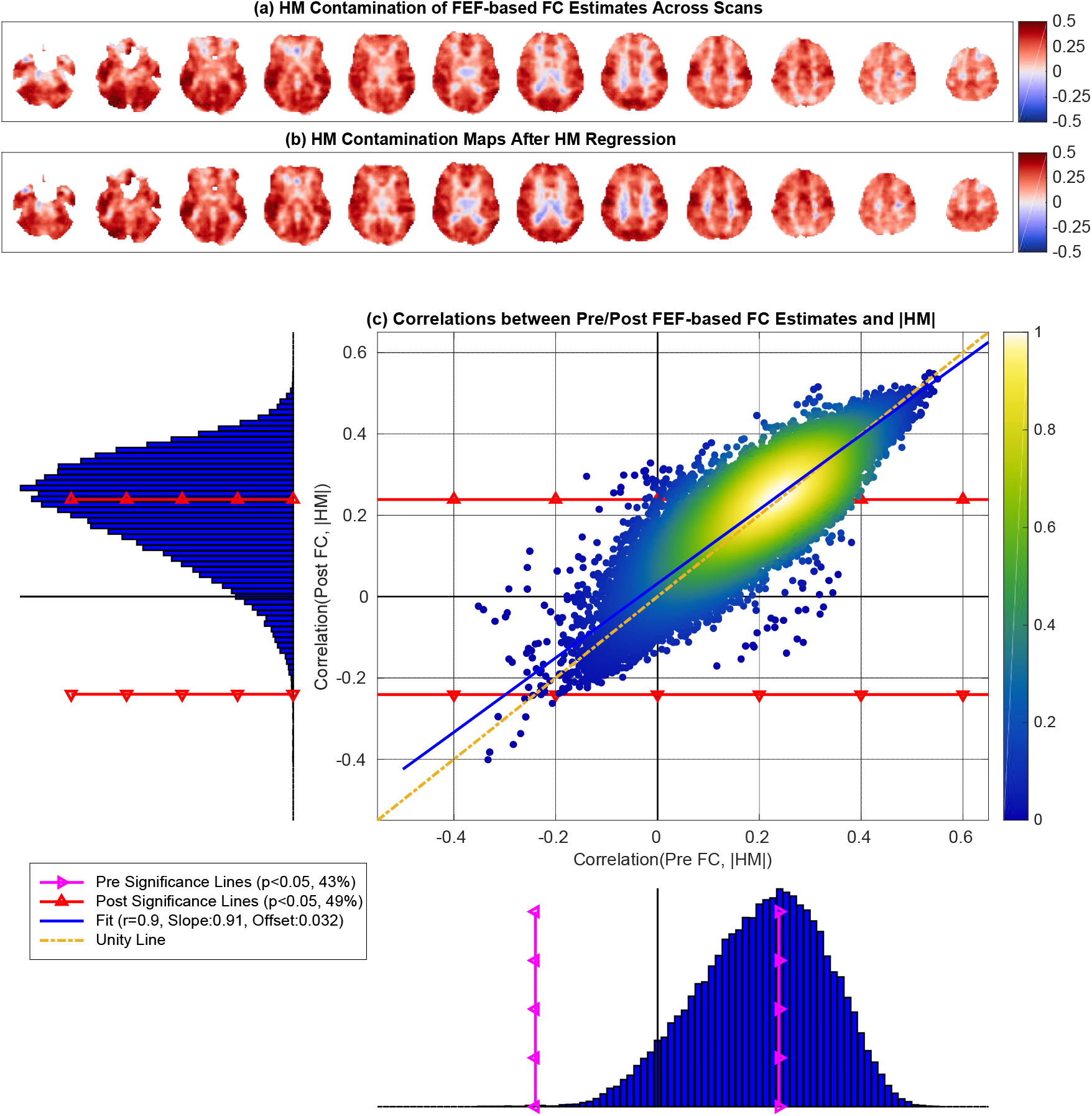
FEF-based HM contamination maps (a) before and (b) after HM regression. These maps were fairly similar to each other (cosine similarity *S* = 0.97) and show widespread correlations between the HM norm and FC estimates across scans. The scatter plot in (c) shows the correlations between the Post FC estimates and HM norm after regression versus the correlations obtained between Pre FC estimates and HM norm. These correlation distributions were significantly related (*r* = 0.9, *p* < 10^−3^) to each other. The linear fit (blue line, Slope= 0.91 and Offset= 0.032) between the two correlation distributions was nearly identical to the line of unity (dashed yellow line). In the bottom histogram, the correlations between the Pre FC estimates and HM norms ranged from *r* = −0.34 to *r* = 0.55 with mean 0.21, and 43% of these correlations were significant (*p* < 0.05). In the sideways histogram on the left, the correlations between the Post FC estimates and HM norms ranged from *r* = −0.40 to *r* = 0.55 with mean 0.22. The post regression significance lines are shown with red lines with triangles. 49% of the Post FC estimates were significantly correlated with HM norms after regression.

**Supplementary Figure 6:**
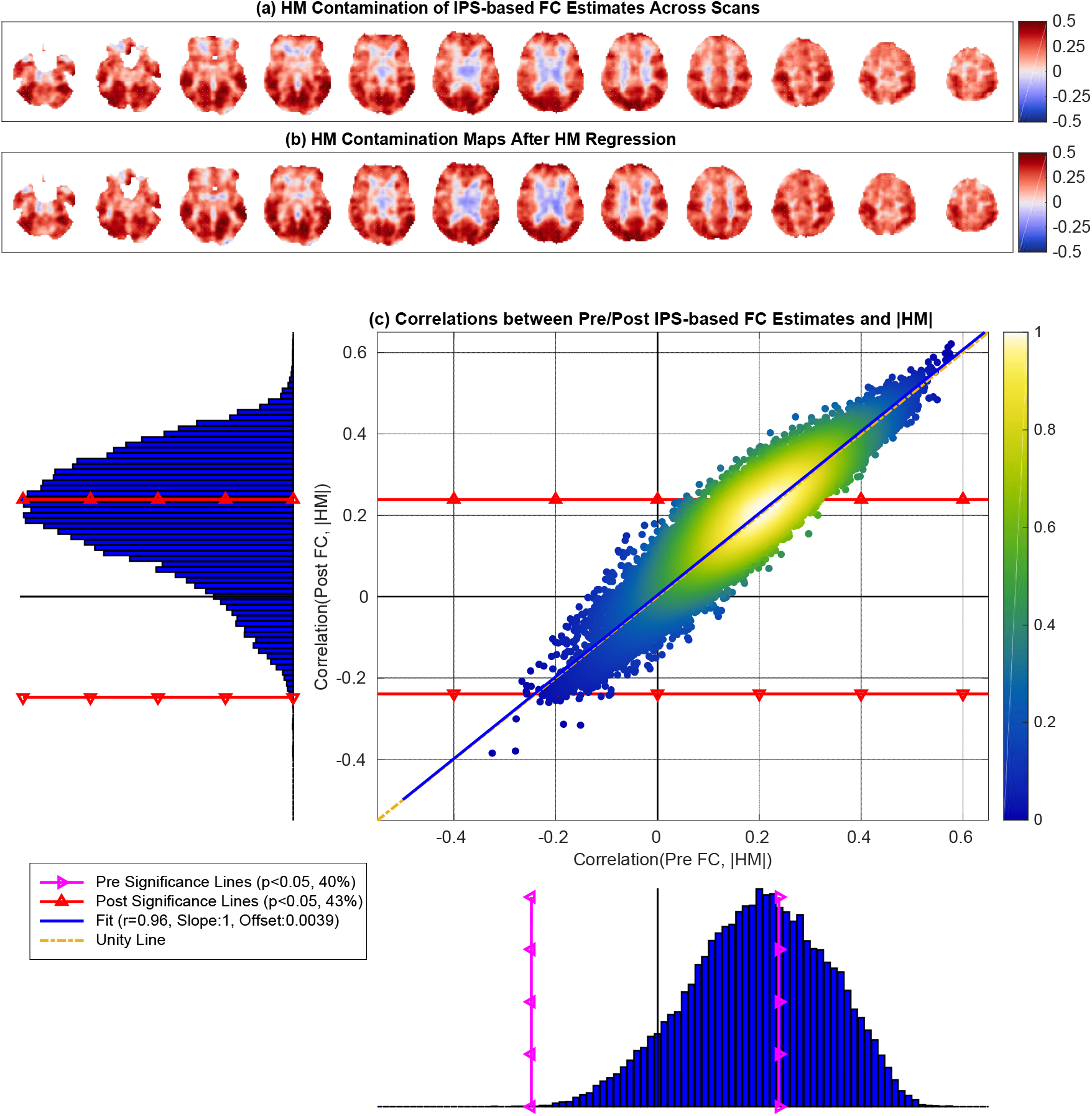
IPS-based HM contamination maps (a) before and (b) after HM regression. These maps were fairly similar to each other (cosine similarity S = 0.99) and show widespread correlations between the HM norm and FC estimates across scans. The scatter plot in (c) shows the correlations between the Post FC estimates and HM norm after regression versus the correlations obtained between Pre FC estimates and HM norm. These correlation distributions were significantly related (*r* = 0.96, *p* < 10^−3^) to each other. The linear fit (blue line, Slope= 1.0 and Offset= 0.0039) between the two correlation distributions was nearly identical to the line of unity (dashed yellow line). In the bottom histogram, the correlations between the Pre FC estimates and HM norms ranged from *r* = −0.33 to *r* = 0.58 with mean 0.20, and 40% of these correlations were significant (*p* < 0.05). In the sideways histogram on the left, the correlations between the Post FC estimates and HM norms ranged from *r* = −0.36 to *r* = 0.62 with mean 0.21. The post regression significance lines are shown with red lines with triangles. 43% of the Post FC estimates were significantly correlated with HM norms after regression.

**Supplementary Figure 7:**
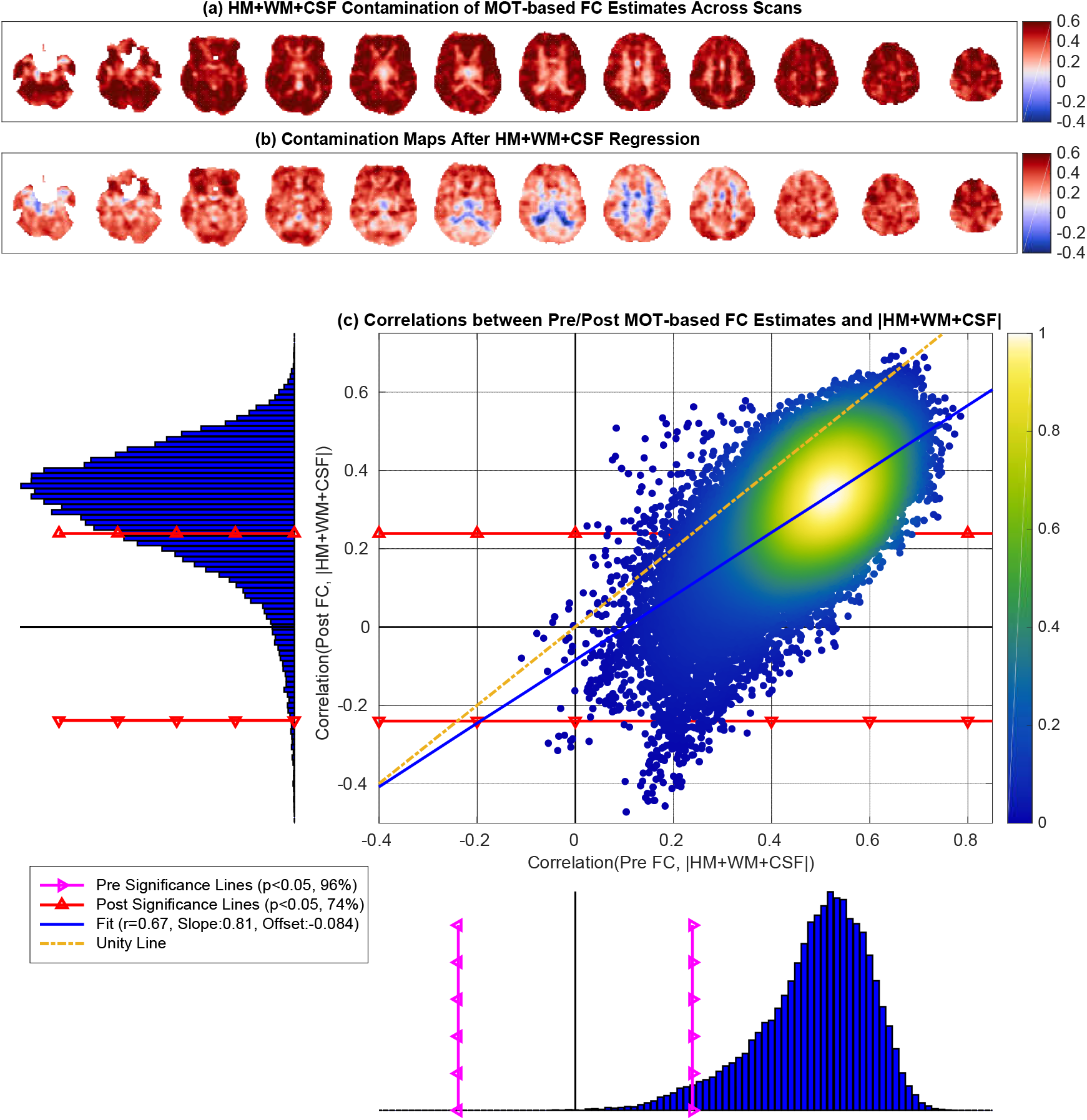
MOT-based nuisance contamination maps (a) before and (b) after HM+WM+CSF regression. There was a visible reduction in the correlation values after regression with a slight increase in anti-correlations with blue regions appearing in (b). In (c) the correlations between the Pre FC estimates and the nuisance norm (x-axis) ranged from *r* = −0.11 to *r* = 0.76 with mean 0.49. The correlations between the Post FC estimates and nuisance norm (y-axis) ranged from *r* = −0.55 to *r* = 0.71 with mean 0.31. There was a strong linear relation between the two correlation distributions (linear fit *r* = 0.67, *p* < 10^−3^). The linear fit between the two correlation distributions was close to the line of unity with a slight reduction in the slope (Slope= 0.81).

**Supplementary Figure 8:**
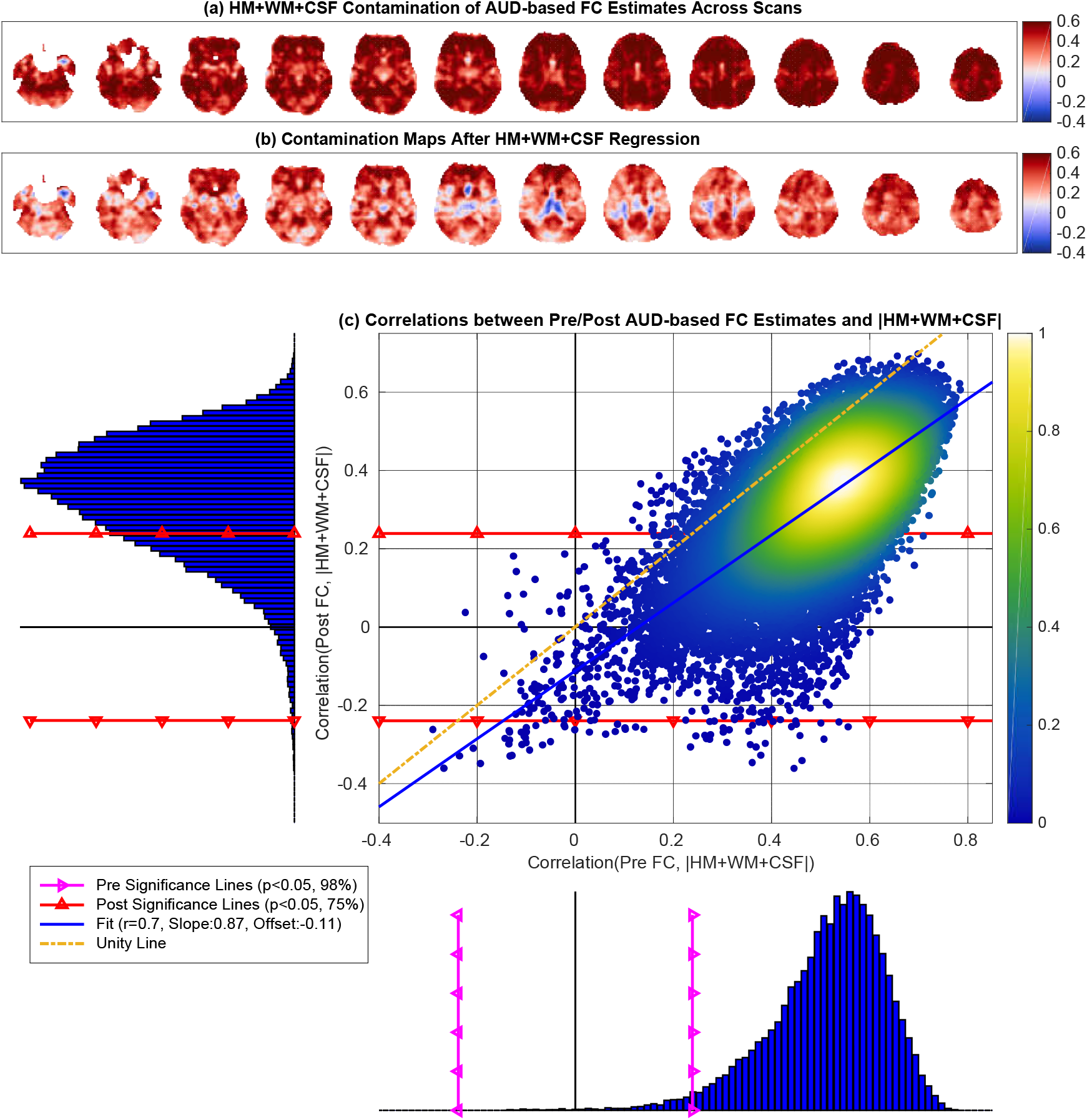
AUD-based nuisance contamination maps (a) before and (b) after HM+WM+CSF regression. In (c) the correlations between the Pre FC estimates and the nuisance norm (x-axis) ranged from *r* = −0.29 to *r* = 0.79 with mean 0.51, and 98% of these correlations were significant (*p* < 0.05). The correlations between the Post FC estimates and nuisance norm (y-axis) ranged from *r* = −0.36 to *r* = 0.70 with mean 0.33, and 74% of these correlations were significant (*p* < 0.05). There was a strong linear relation between the two correlation distributions (linear fit *r* = 0.70, *p* < 10^−3^). The linear fit between the two correlation distributions was close to the line of unity with a slight reduction in the slope (Slope= 0.87) and a small negative offset of −0.11.

**Supplementary Figure 9:**
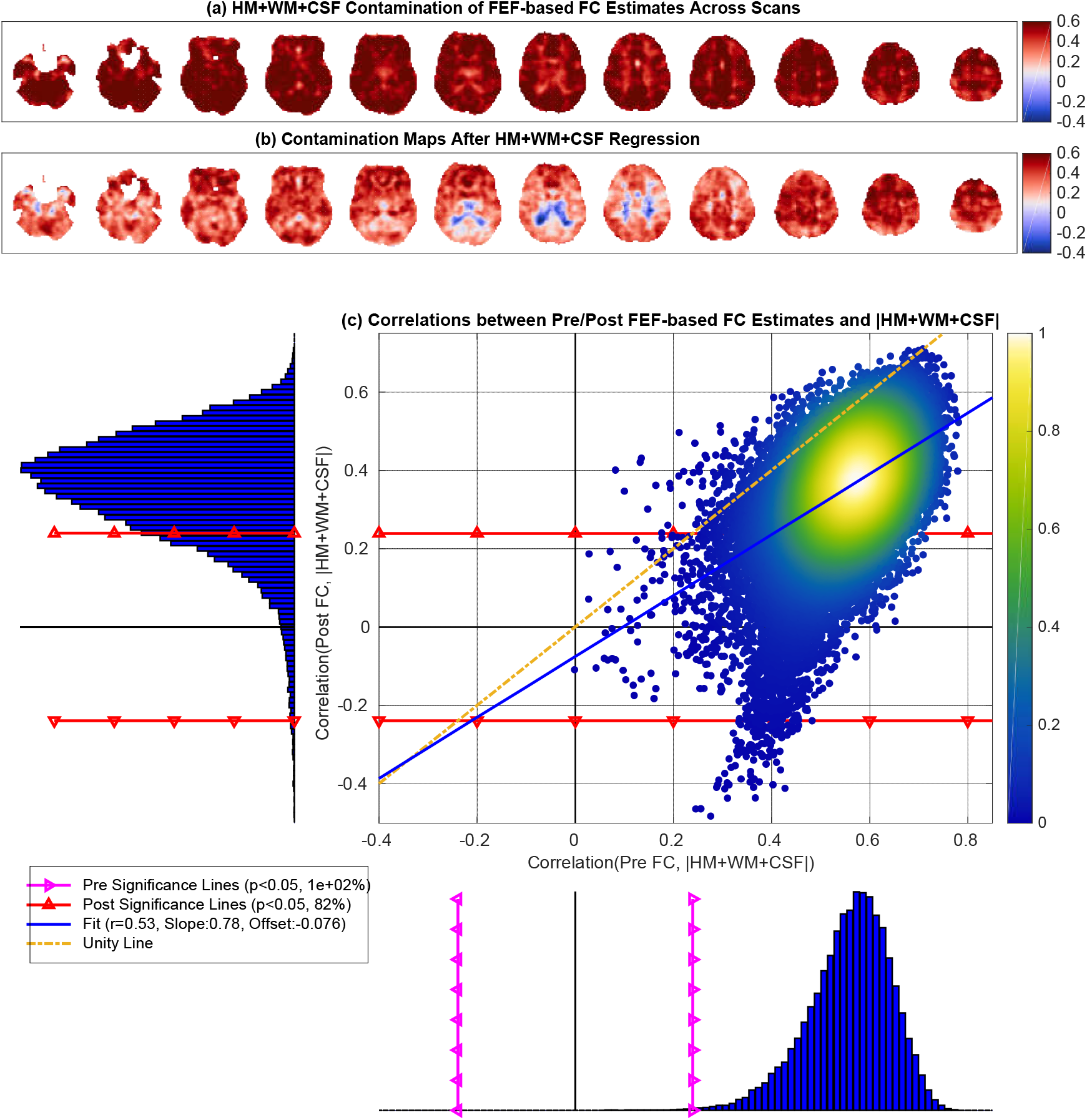
FEF-based nuisance contamination maps (a) before and (b) after HM+WM+CSF regression. In (c) the correlations between the Pre FC estimates and the nuisance norm (x-axis) ranged from *r* = 0.0 to *r* = 0.78 with mean 0.56, and 98% of these correlations were significant (*p* < 0.05). The correlations between the Post FC estimates and nuisance norm (y-axis) ranged from *r* = −0.51 to *r* = 0.71 with mean 0.35, and 75% of these correlations were significant (*p* < 0.05). There was a strong linear relation between the two correlation distributions (linear fit *r* = 0.53, *p* < 10^−3^). The linear fit between the two correlation distributions was close to the line of unity with a slight reduction in the slope (Slope= 0.78) and a small negative offset of −0.08.

**Supplementary Figure 10:**
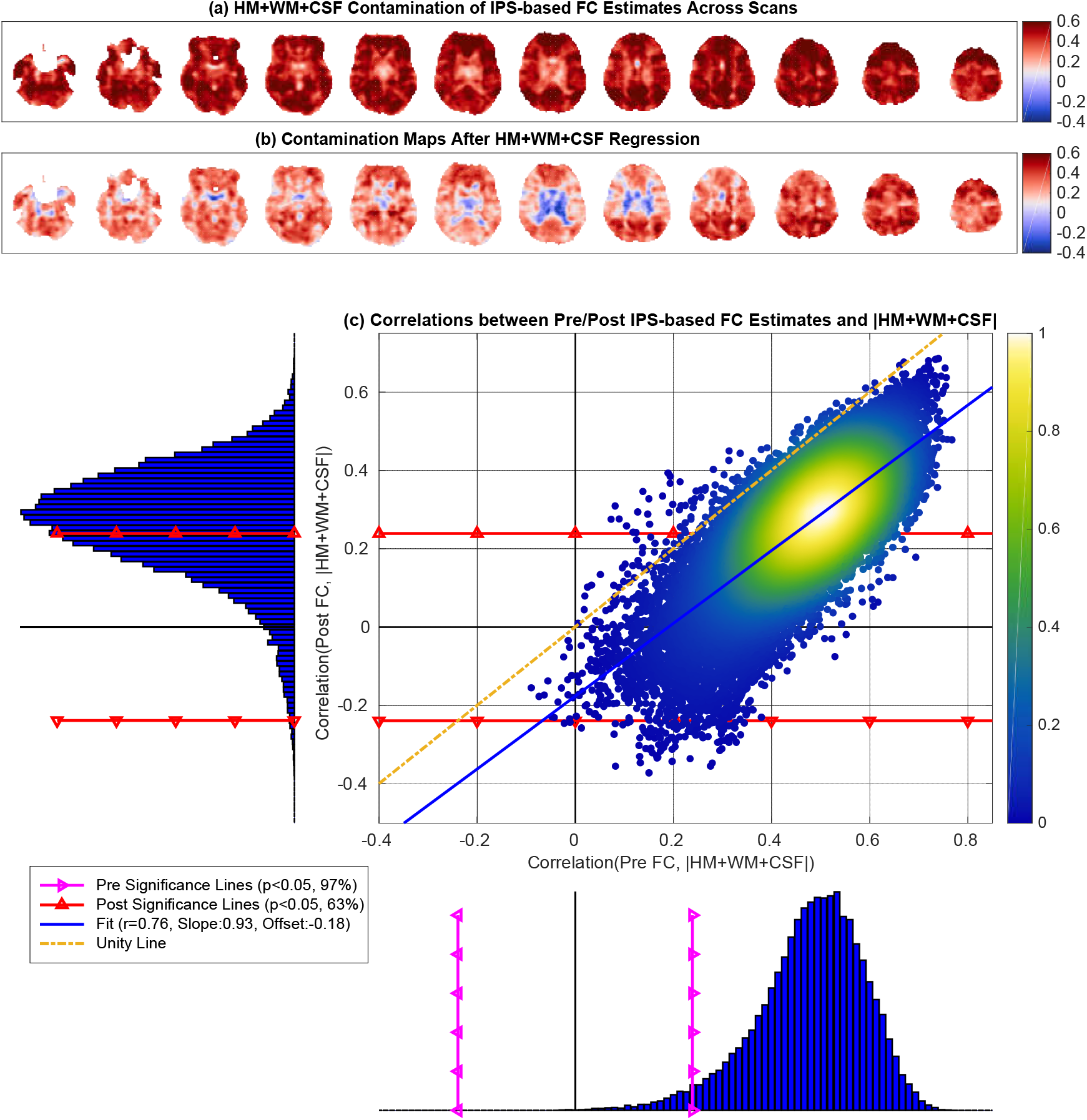
IPS-based nuisance contamination maps (a) before and (b) after HM+WM+CSF regression. In (c) the correlations between the Pre FC estimates and the nuisance norm (x-axis) ranged from *r* = −0.1 to *r* = 0.77 with mean 0.48, and 97% of these correlations were significant (*p* < 0.05). The correlations between the Post FC estimates and nuisance norm (y-axis) ranged from *r* = −0.37 to *r* = 0.68 with mean 0.27, and 63% of these correlations were significant (*p* < 0.05). There was a strong linear relation between the two correlation distributions (linear fit *r* = 0.76, *p* < 10^−3^). The linear fit between the two correlation distributions was close to the line of unity with a slight reduction in the slope (Slope= 0.93) and a small negative offset of −0.18.

**Supplementary Figure 11:**
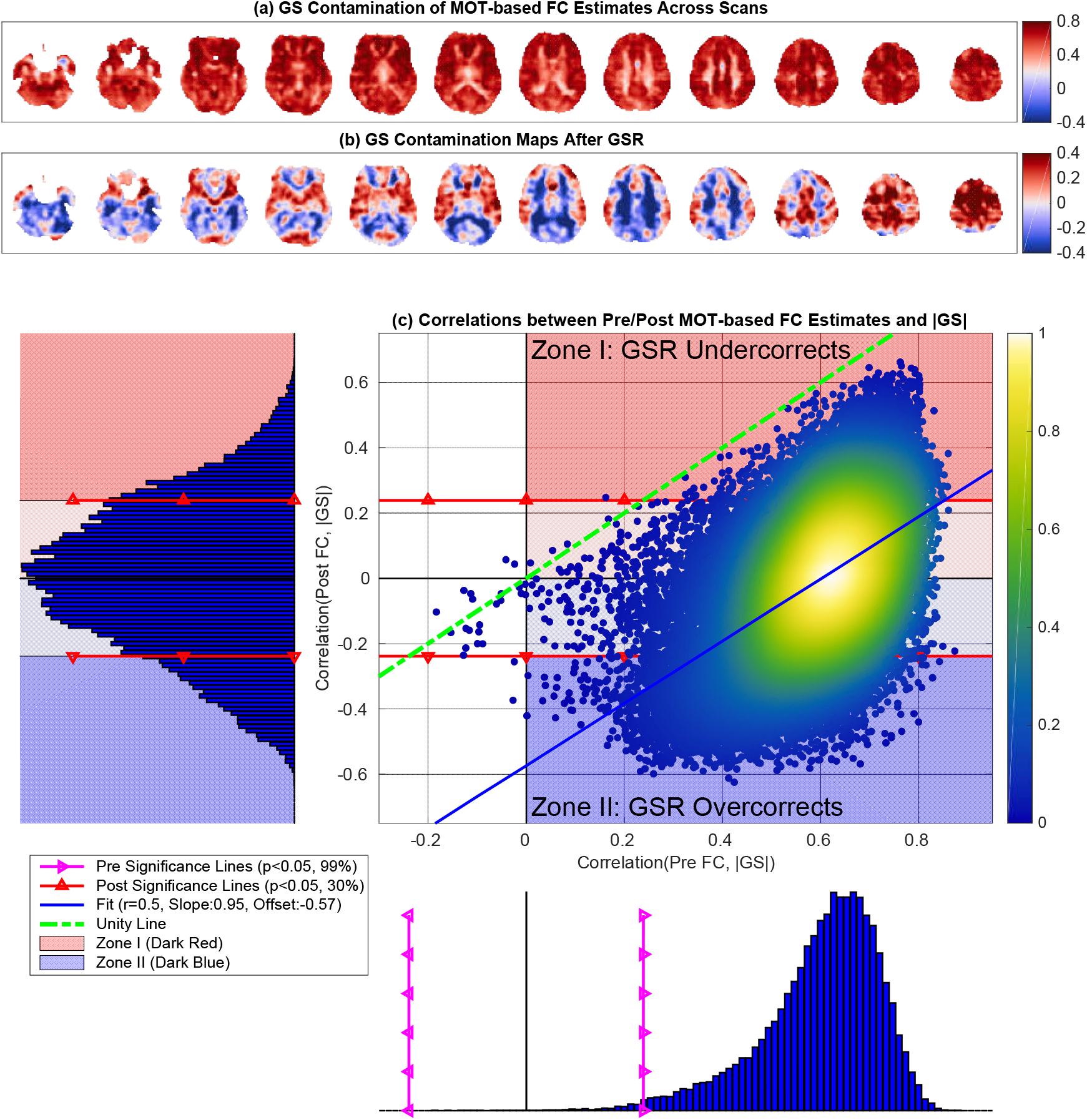
MOT-based GS contamination maps obtained both (a) before and (b) after GSR. In panel (a) the contamination map exhibits predominantly positive correlations (99%) between the Pre FC estimates and GS norms. In (b) the contamination map after GSR exhibits both positive (50%) and negative (50%) correlations between the GS norms and Post FC estimates. In (c) the correlations between the Pre FC estimates and the nuisance norm (x-axis) ranged from *r* = −0.18 to *r* = 0.87 with mean 0.60, and 99% of these correlations were significant (*p* < 0.05). The correlations between the Post FC estimates and nuisance norm (y-axis) ranged from *r* = −0.62 to *r* = 0.66 with mean 0.0, and 30% of these correlations were significant (*p* < 0.05). There was a strong linear relation between the two correlation distributions (linear fit *r* = 0.50, *p* < 10^−3^), and the linear fit (blue line) was fairly parallel to the line of unity with slope of 0.95 and a very large negative offset of −0.57.

**Supplementary Figure 12:**
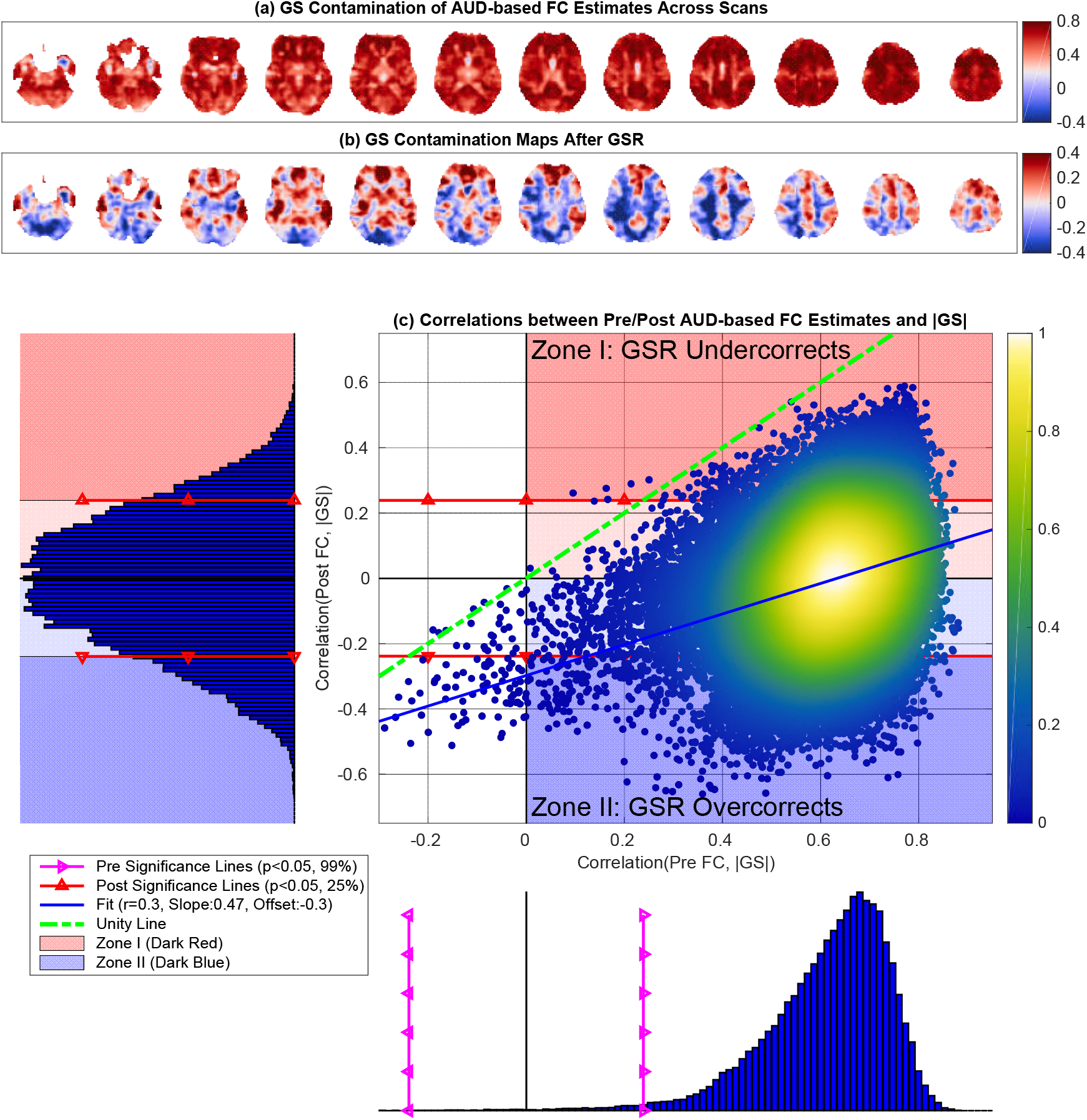
AUD-based GS contamination maps obtained both (a) before and (b) after GSR. In panel (a) the contamination map exhibits predominantly positive correlations (99%) between the Pre FC estimates and GS norms. In (b) the contamination map after GSR exhibits both positive (48%) and negative (52%) correlations between the GS norms and Post FC estimates. In (c) the correlations between the Pre FC estimates and the nuisance norm (x-axis) ranged from *r* = −0.35 to *r* = 0.88 with mean 0.61, and 99% of these correlations were significant (*p* < 0.05). The correlations between the Post FC estimates and nuisance norm (y-axis) ranged from *r* = −0.66 to *r* = 0.59 with mean 0.01, and 25% of these correlations were significant (*p* < 0.05). There was a linear relation between the two correlation distributions (linear fit *r* = 0.30, *p* < 10^−3^), and the linear fit (blue line) was fairly parallel to the line of unity with slope of 0.47 and a large negative offset of −0.30.

**Supplementary Figure 13:**
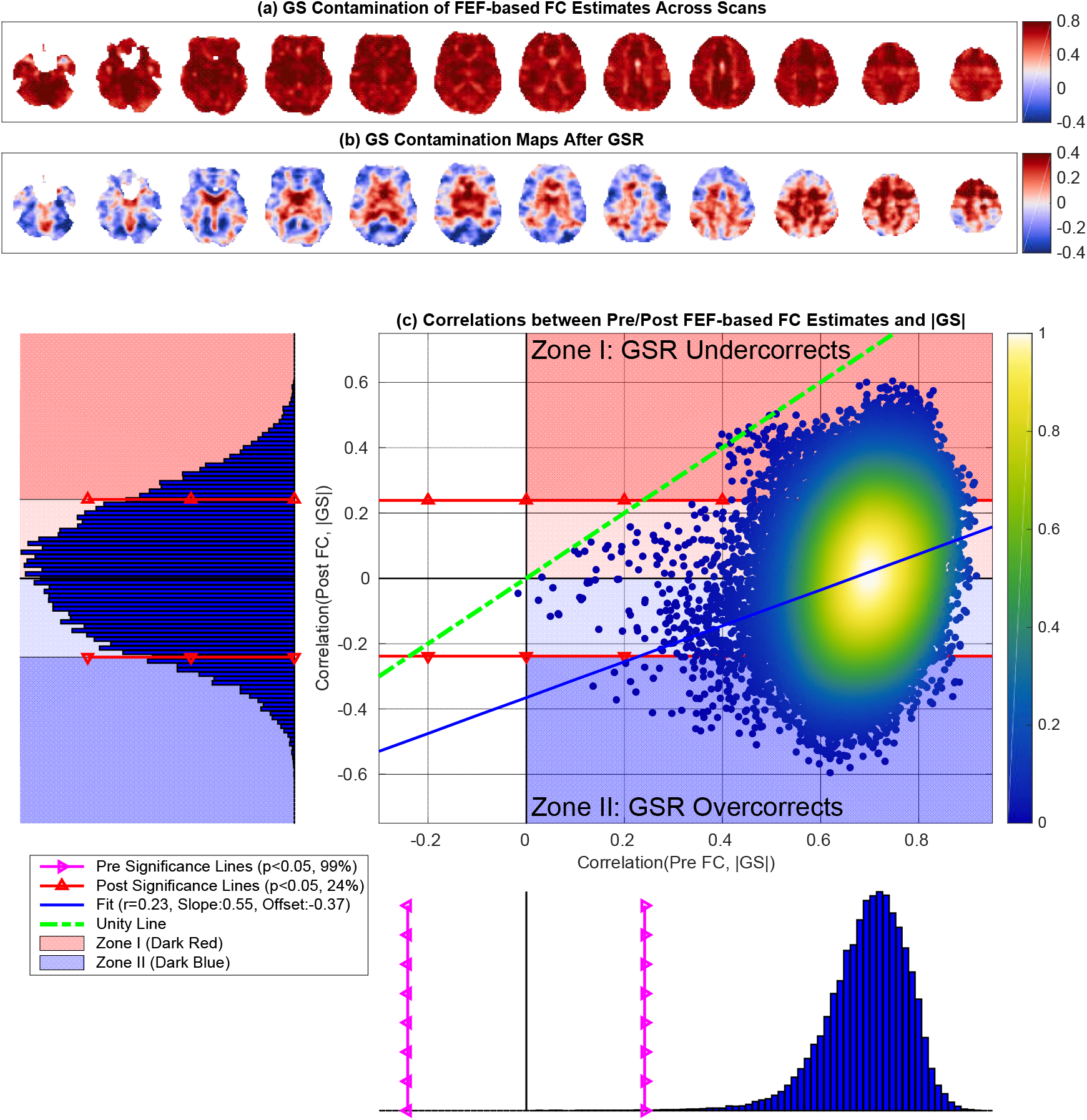
FEF-based GS contamination maps obtained both (a) before and (b) after GSR. In panel (a) the contamination map exhibits predominantly positive correlations (100%) between Pre FC estimates and GS norm. In (b) the contamination map after GSR exhibits both positive (53%) and negative (48%) correlations between the GS norms and Post FC estimates. In (c) the correlations between the Pre FC estimates and the nuisance norm (x-axis) ranged from *r* = −0.02 to *r* = 0.92 with mean 0.70, and 100% of these correlations were significant (*p* < 0.05). The correlations between the Post FC estimates and nuisance norm (y-axis) ranged from *r* = −0.60 to *r* = 0.60 with mean 0, and 24% of these correlations were significant (*p* < 0.05). There was a weak linear relation between the two correlation distributions (linear fit *r* = 0.23, *p* < 10^−3^), and the linear fit (blue line) was moderately parallel to the line of unity with a slope of 0.55 and a large negative offsetof −0.37.

**Supplementary Figure 14:**
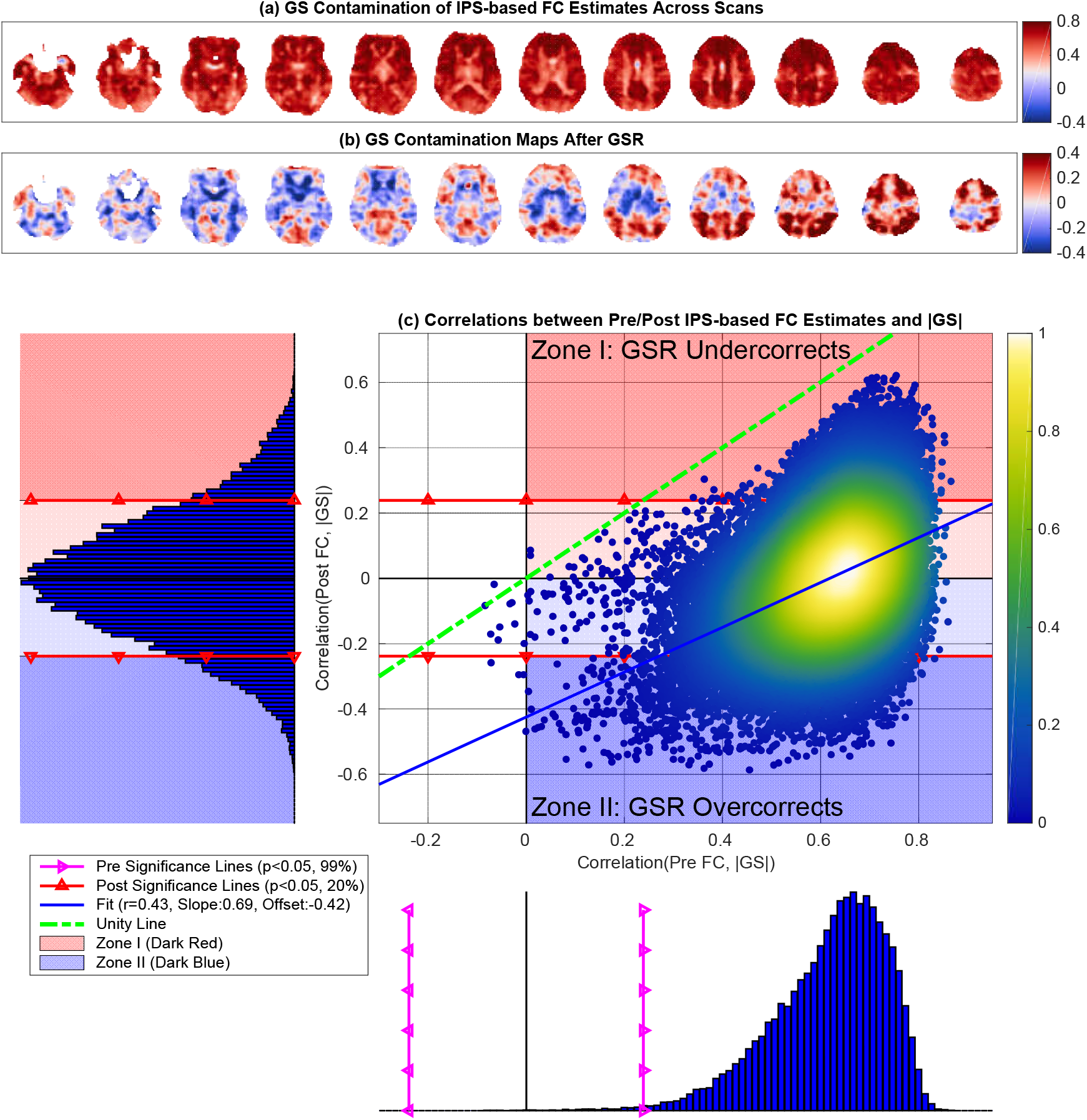
IPS-based GS contamination maps obtained both (a) before and (b) after GSR. In panel (a) the contamination map exhibits predominantly positive correlations (100%) between Pre FC estimates and GS norm. In (b) the contamination map after GSR exhibits both positive (49%) and negative (51%) correlations between the GS norms and Post FC estimates. In (c) the correlations between the Pre FC estimates and the nuisance norm (x-axis) ranged from *r* = −0.1 to *r* = 0.87 with mean 0.62, and 100% of these correlations were significant (*p* < 0.05). The correlations between the Post FC estimates and nuisance norm (y-axis) ranged from *r* = −0.59 to *r* = 0.62 with mean 0, and 20% of these correlations were significant (*p* < 0.05). There was a linear relation between the two correlation distributions (linear fit *r* = 0.43, *p* < 10^−3^), and the linear fit (blue line) was fairly parallel to the line of unity with a slope of 0.69 and a large negative offset of −0.42.

**Supplementary Figure 15:**
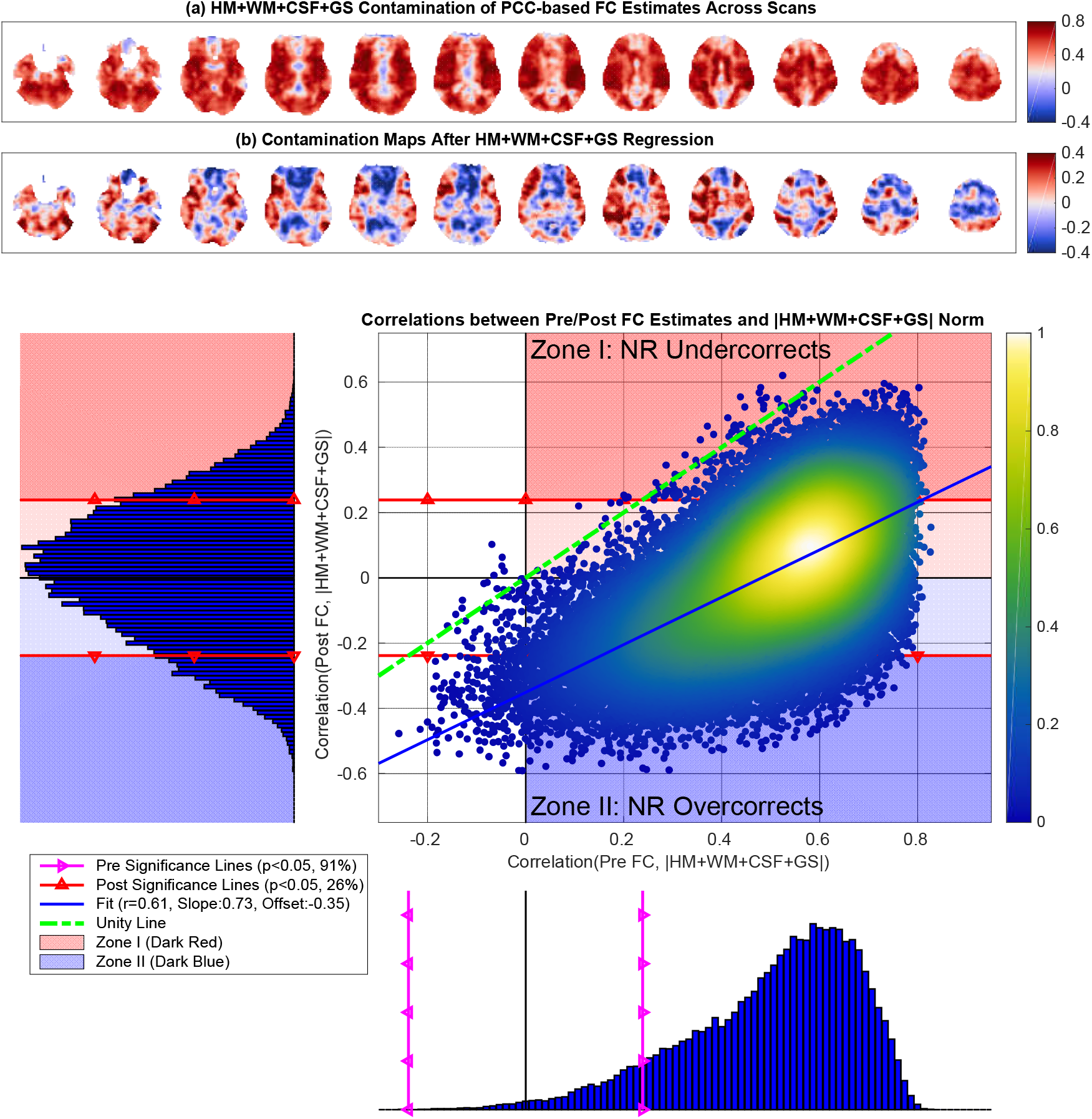
PCC-based HM+WM+CSF+GS contamination maps obtained both (a) before and (b) after multiple regression. In panel (a) the contamination map mostly exhibits predominantly positive correlations (99%) between Pre FC estimates and nuisance norm. In (b) the contamination map after nuisance regression exhibits both positive (54%) and negative (46%) correlations between the nuisance norm and Post FC estimates. In (c) the correlations between the Pre FC estimates and the nuisance norm (x-axis) ranged from *r* = −0.26 to *r* = 0.83 with mean 0.50, and 99% of these correlations were significant (*p* < 0.05). The correlations between the Post FC estimates and nuisance norm (y-axis) ranged from *r* = −0.59 to *r* = 0.62 with mean 0.01, and 26% of these correlations were significant (*p* < 0.05). There was a linear relation between the two correlation distributions (linear fit *r* = 0.61, *p* < 10^−3^), and the linear fit (blue line) was fairly parallel to the line of unity with a slope of 0.73 and a large negative offset of −0.35. These results were very similar to those obtained when performing GSR alone.

